# Establishing CRISPR-Cas9 in the sexually dimorphic moss, *Ceratodon purpureus*

**DOI:** 10.1101/2023.10.25.562971

**Authors:** Emilie-Katherine Tavernier, Emily Lockwood, Pierre-François Perroud, Fabien Nogué, Stuart F. McDaniel

## Abstract

The development of CRISPR technologies provides a powerful tool for understanding the evolution and functionality of essential biological processes. Here we demonstrate successful CRISPR-Cas9 genome editing in the dioecious moss species, *Ceratodon purpureus*. Using an existing selection system from the distantly related hermaphroditic moss, *Physcomitrium patens*, we generated knock-outs of the *APT* reporter gene by employing CRISPR targeted mutagenesis under expression of native U6 snRNA promoters. Next, we used the native homology-directed repair (HDR) pathway, combined with CRISPR-Cas9, to knock-in two reporter genes under expression of an endogenous RPS5A promoter in a newly developed landing site in *C. purpureus*. Our results show that the molecular tools developed in *P. patens* can be extended to other mosses across this ecologically important and developmentally variable group. These findings pave the way for precise and powerful experiments aimed at identifying the genetic basis of key functional variation within the bryophytes and between the bryophytes and other land plants.

**Significance Statement:** We have developed CRISPR-Cas9 genome editing tools and protocols for the sexually dimorphic moss, *Ceratodon purpureus* to generate gene knock-outs and knock-ins within targeted loci. This work facilitates future functional genomic experiments in this fast growing, haploid, dioecious system and suggests these tools will function across much of moss diversity.

## Introduction

The application of clustered regularly interspaced short palindromic repeats (CRISPR)-associated protein (Cas) gene editing technology has increased the speed and precision of gene functional analyses (Zhang et al., 2021). CRISPR-Cas9 is an RNA-protein complex that is engineered to generate genomic double stranded DNA breaks (DSBs) at the specified location of the user and is widely used to generate targeted gene knock-outs and, more rarely, gene knock-ins within living cells (Knott and Doudna, 2018). Part of the power of CRISPR-Cas is that it can be employed in species with important biological attributes that lack the genetic resources developed for major model systems (Shan et al., 2020).

CRISPR-Cas genetic engineering is well developed in the monoecious model moss, *Physcomitrium patens* (Rensing et al., 2020). In addition to gene disruption, *P. patens* readily accommodates gene targeting using the homology directed repair (HDR) pathway with CRISPR-Cas9 to knock-in genes into selected genomic regions (Collonnier et al., 2017b; Mara et al., 2019). HDR guided gene insertion uses a landsite for ectopic transgene expression, that is in a genomic region that is actively transcribed but does not disrupt another gene (Mallett et al., 2019). However, *P. patens* is not representative of the diversity of mosses, most of which are dioecious. The sex determination system in dioecious mosses is regulated by a UV chromosome system, in which female gametophytes possess a U and males possess a V. The bryophyte UV sex chromosome system is conserved across peristomate mosses and may be shared by the liverwort *Marchantia polymorpha* (Carey et al., 2021). The antiquity of dioecy in this system suggests that the sex chromosomes may harbor genes that control sex-limited processes, providing a strong justification for developing gene targeting tools in a dioecious moss.

The dioecious moss, *Ceratodon purpureus*, is an obvious candidate in which to develop gene targeting tools. *C. purpureus* has long been used as a model system in developmental biology (Cove and Quatrano, 2006; Kern and Sack, 1999; Mittmann et al., 2009; Thornton et al., 2005) and ecological genetics studies (Jules and Shaw, 1994; McDaniel, 2005; McDaniel et al., 2008, 2007; Nieto-Lugilde et al., 2018a, 2018b). The species is easy to cultivate in laboratory conditions, amenable to chemical or ultraviolet mutagenesis, and natural populations are highly polymorphic and common in disturbed areas on all continents (McDaniel and Shaw, 2005). *C. purpureus* has only experienced one whole genome duplication, approximately 70mya (Szövényi et al., 2015) unlike the two genome duplications evident in the *P. patens* genome (Rensing et al., 2020, 2007). A single duplication means gene-family sizes, on average, are about half the size in *C. purpureus* than those in *P. patens*. Although gene targeting via HDR has been demonstrated in this species, *C. purpureus* lacks a landing site comparable to that of *P. patens* (Kamisugi et al., 2006; Trouiller et al., 2007), as well as protein promoters that can drive the transcription of genes of interest (Horstmann et al., 2004).

Translating CRISPR-based forward genetic studies requires vetting several potentially species-specific protocols and molecular components in the target organism. To streamline troubleshooting in plants, we can harness the endogenous adenine-phosphoribosyl transferase (*APT*) gene expressed by algae, mosses, and many flowering plants to efficiently select knock-out transformants (Collonnier et al., 2017a; Guzmán-Zapata et al., 2019; Trouiller et al., 2006). As a phosphoribosyl transferase, the APT enzyme catalyzes the conversion of free adenines to their nucleotide form—adenosine monophosphate (AMP). Consequently, APT will also metabolize the toxic adenine analog, 2-fluoroadenine (2-FA), producing the lethal 2-fluoro-AMP, which results in the death of the plant. Therefore, knocking out *APT* confers resistance to 2-FA since the plants can no longer generate AMP analogs with this pathway (Gaillard et al., 1998). The 2-FA resistant phenotype reduces the time necessary to screen transformants, making this system a powerful tool in troubleshooting gene editing.

Here, we test several components of the CRISPR-Cas9 gene editing system in *C. purpureus* using the endogenous *APT* knock-out selection method. We then demonstrate the use of CRISPR-Cas9 nuclease activity in conjunction with homology-based gene-targeting to insert two fluorophores, GFP and mCherry, in a newly established landing site in *C. purpureus* under the regulation of a native *C. purpureus* RPS5a promoter. The combination of experimental tractability, modest gene-family sizes, and abundant polymorphism, including sexual dimorphism, provides a strong justification for developing CRISPR genome editing tools in *C. purpureus*. These results provide a clear path toward functional genomic studies using knock-outs and knock-ins in the emerging model, *C. purpureus*, and showcase the potential to incorporate powerful transgenic tools into bryophyte comparative biology.

## Results

### *Efficient gene knock-out in* C. purpureus *with RNA-guided Cas9 nuclease*

To investigate whether CRISPR-Cas could be used for genome editing in *C. purpureus*, we chose to target the *APT* (adenine phosphoribosyl transferase) gene. Knocking out *APT* renders the plant unable to metabolize 2-FA, and consequently confers resistance to the toxin. We identified and characterized *CpAPT* (CepurGG1.12G128200; https://phytozome-next.jgi.doe.gov/) by searching the *C. purpureus* R40 and GG1 genomes v1.1 (https://phytozome-next.jgi.doe.gov/) using the previously identified partial *APT* gene sequence (DQ265803) as a Basic Local Alignment Search Tool query. To knock-out the *C. purpureus APT* (Figure 1) we designed two sgRNAs, CpAPT1 and CpAPT2.

**Figure 1.**
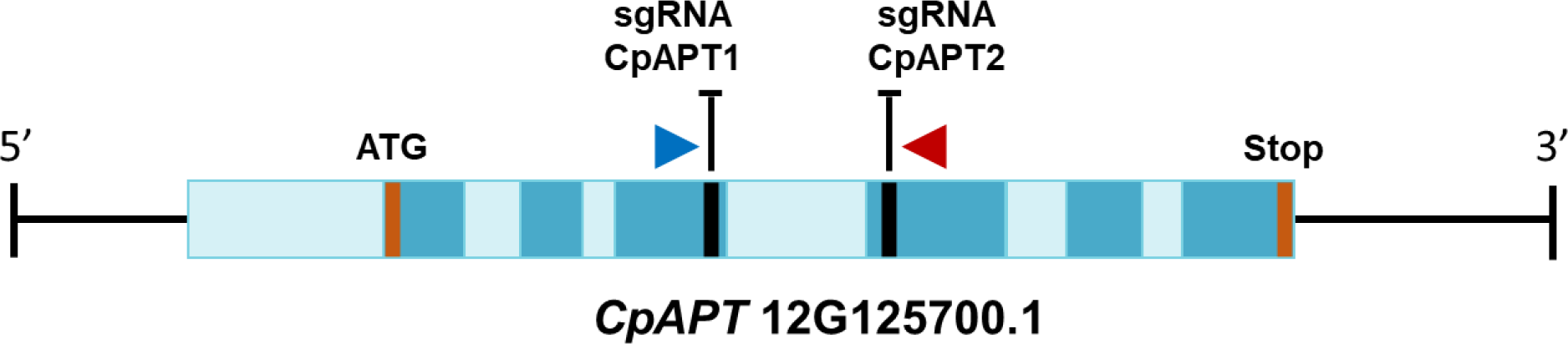
Scheme of *C. purpureus* 12G125700.1 adenine phosphoribosyl transferase (*APT*) gene with sgRNA 20bp targets and primer binding sites marked. 20bp target sites are indicated in black: CpAPT1 and CpAPT2. Forward primers (blue) and reverse primers (red) show which regions were amplified with PCR and sequenced. Light blue boxes: introns. Dark blue boxes: exons.

For expression of the sgRNAs, we identified the promoters of four different *C. purpureus* snRNAs, taken from chromosomes 7, 9, 10 and 12 (Figure 2, Table S4). We used jBrowse log-scale RNA-seq coverage data to confirm that each snRNA gene was transcribed (Figure S3). We defined the promotor as the 315bp directly upstream of the gene to ensure it contained the Upstream Sequence Element (USE) as well as the TATA box (Filipowicz et al. 1990).

**Figure 2.**
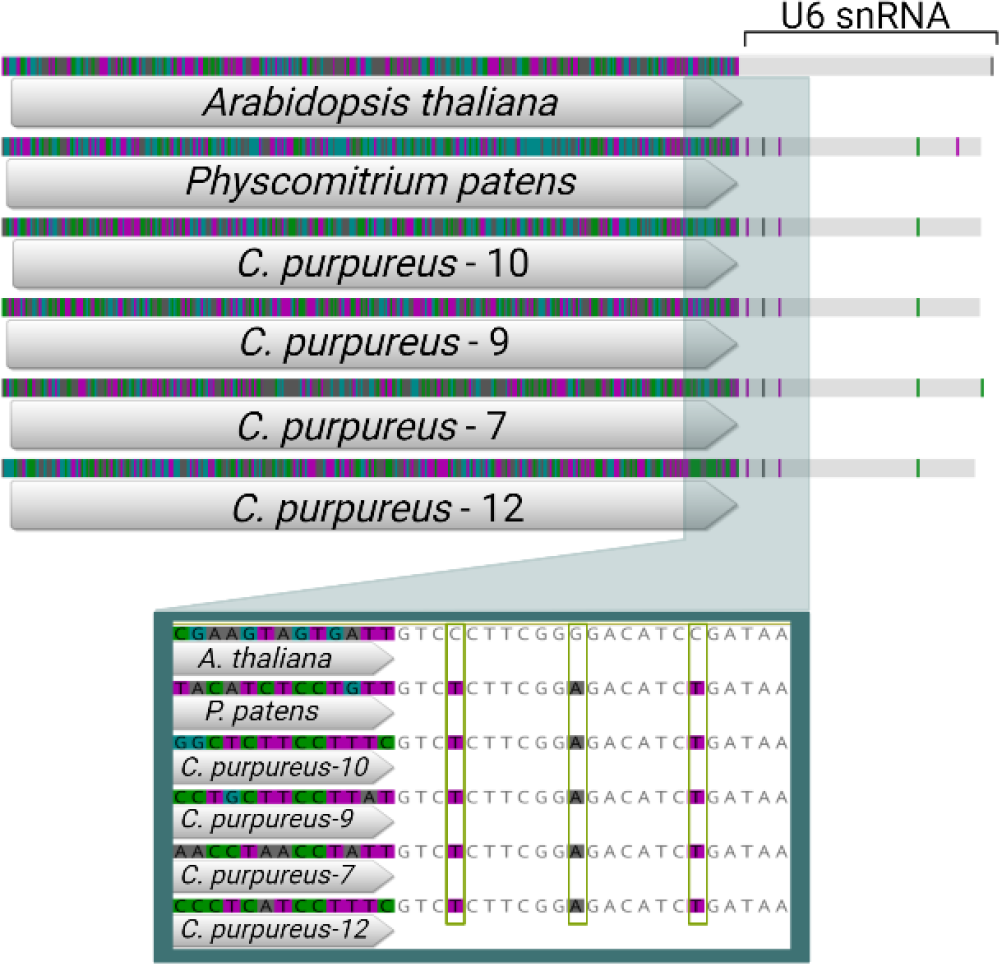
Alignment of all highly variable U6 promoters used in this study, highlighting the SNPs between the *Arabidopsis thaliana* snRNA and the *P. patens* and *C. purpureus* conserved U6 snRNA sequences.

We transformed *C. purpureus* protoplasts with the two sgRNAs, sgRNA_CpAPT1 and sgRNA_CpAPT2, driven by one of the four *C. purpureus* U6 promoters or by the established *P. patens* U6 promoter (Collonnier et al., 2017), in tandem with pAct-Cas9, via PEG-mediated transfection. All 96 2-FA resistant *CpAPT*-KO plants that we sanger sequenced had mutations at one or both *CpAPT* cut sites. These data demonstrate that CRISPR-Cas9 is functional in *C. purpureus* and that all U6 promoters successfully recruited the RNA polymerase III to transcribe the sgRNAs (Figure 3). The mutation rates (expressed in percentages) were estimated by dividing the number of 2-FA-resistant plants by the number of regenerating plants observed just before the transfer to 2-FA. The mutation rates obtained with the different U6 promoters varied between promoters, with the male isolate R40 consistently producing more mutants from each *APT-KO* construct over the female, 15-12-12 (Figure 3, Table 1). We found no correlation between the native expression levels of snRNA using jBrowse (https://phytozome-next.jgi.doe.gov/jbrowse/), and the transcriptional capability of individual U6 promoters to transcribe each sgRNA (Figure S3).

**Table 1.** Counts of protoplast regeneration efficiency and gene editing efficiency for *C. purpureus* APT-KO mutants using each of the evaluated U6 promoters.

| U6 Promoter | Protoplasts regenerated |  | Protoplast regeneration efficiency |  | 2-FA+ plants |  | Gene editing efficiency |  |
| --- | --- | --- | --- | --- | --- | --- | --- | --- |
| Cp7 | 24276 | 17526 | 0.1618 | 0.1168 | 106 | 134 | 0.436 | 0.7646 |
| Cp9 | 13538 | 25381 | 0.0903 | 0.1692 | 36 | 123 | 0.265 | 0.4846 |
| Cp10 | 16153 | 28122 | 0.1077 | 0.1875 | 114 | 219 | 0.705 | 0.7788 |
| Cp12 | 13611 | 13421 | 0.0907 | 0.0895 | 2 | 55 | 0.014 | 0.4098 |
| Pp | 11916 | 28691 | 0.0794 | 0.1913 | 41 | 111 | 0.344 | 0.3869 |

15-12-12 is boxed in gray. R40 is boxed in white.
Gene editing (GE) represents the frequency of 2-FA resistant plants among the regenerated protoplasts

**Figure 3.**
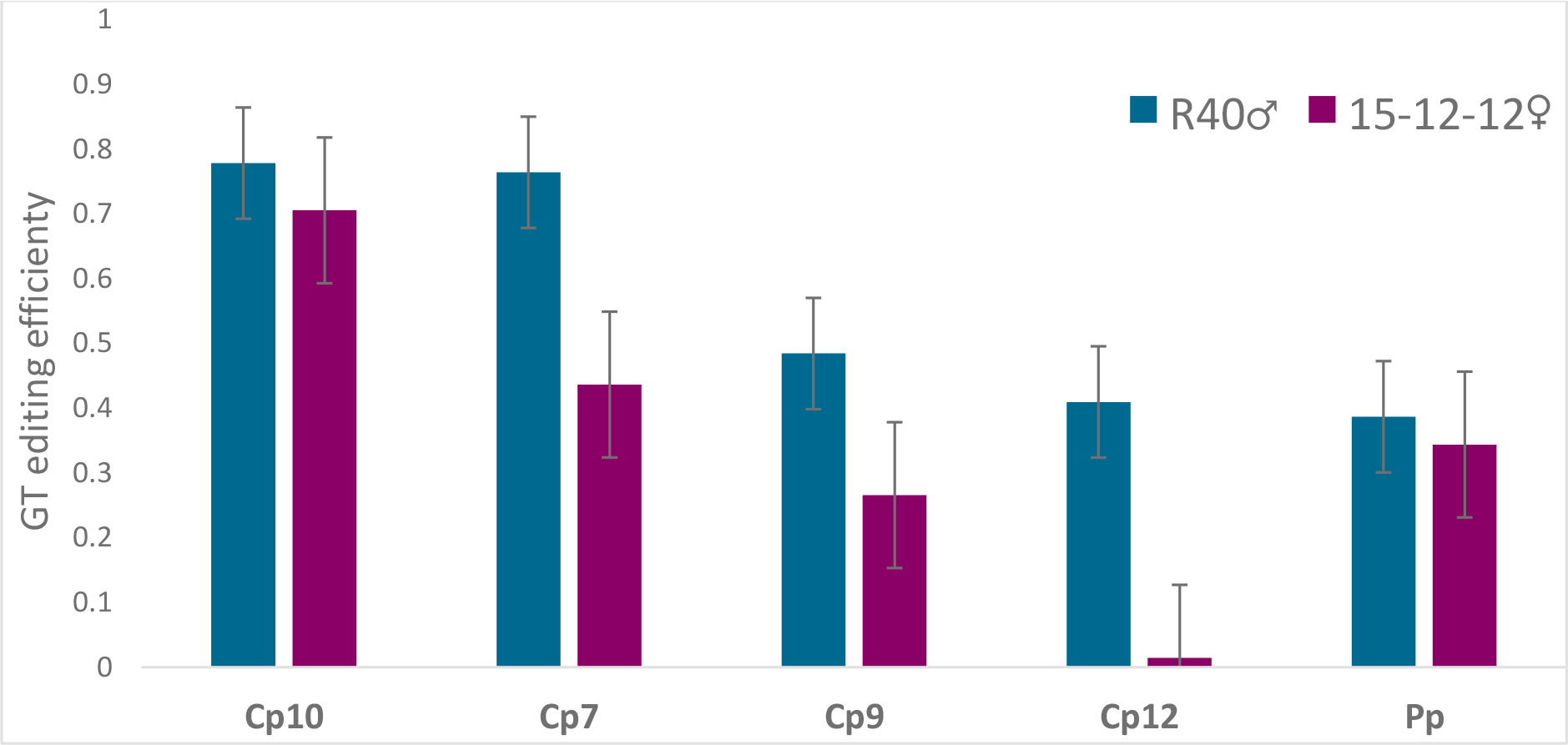
Relative Gene Editing (GE) efficiency of sgRNA_CpAPT1 and sgRNA_CpAPT2 vectors constructed with *C. purpureus* U6 promoters from autosomes 7, 9 10 and 12, and *P. patens* U6 promoter from autosome 1. Approximately 150,000 protoplasts were transformed per reaction.

### *Cas9 nuclease double strand breaks can be repaired by both cNHEJ and Alt-EJ in* C. purpureus

The efficiency between sgRNA_CpAPT1 and sgRNA_CpAPT2 differed between each *C. purpureus* genotype. In female isolates 15-12-12 and B150, CpAPT1 cut 100% of the time, and CpAPT 2 cut 28% of the time. In male isolates R40 and B190, CpAPT1 cut 89% of the time, while CpAPT2 cut 58% of the time (Figure S4).

To repair DSBs, *C. purpureus* used both the cNHEJ and Alt-EJ pathways. 52 out of 100 genotyped *CpAPT-*KO plants displayed a mixed sequencing chromatogram signal at a sgRNA cut site. We considered them as mosaics and eliminated them from further analyses. The remaining 48 *CpAPT-*KO plants were suitable to distinguish between MMEJ or cNHEJ repair pathway. The MMEJ pathway repairs DSBs using small microhomologies of two to seven base pairs, leaving clean repairs. The cNHEJ pathway forgoes the need for homology arms by direct ligation between the DNA strands and can lead to more random insertions and deletions in the genome. Both DNA repair mechanisms occur during the cell cycle (Sakamoto et al., 2021). These data show that the *C. purpureus* male and female isolates we tested demonstrate the same ratio of MMEJ:NHEJ repairs, and that three of the four isolates selectively use the MMEJ pathway over cNHEJ (Figure 4).

**Figure 4.**
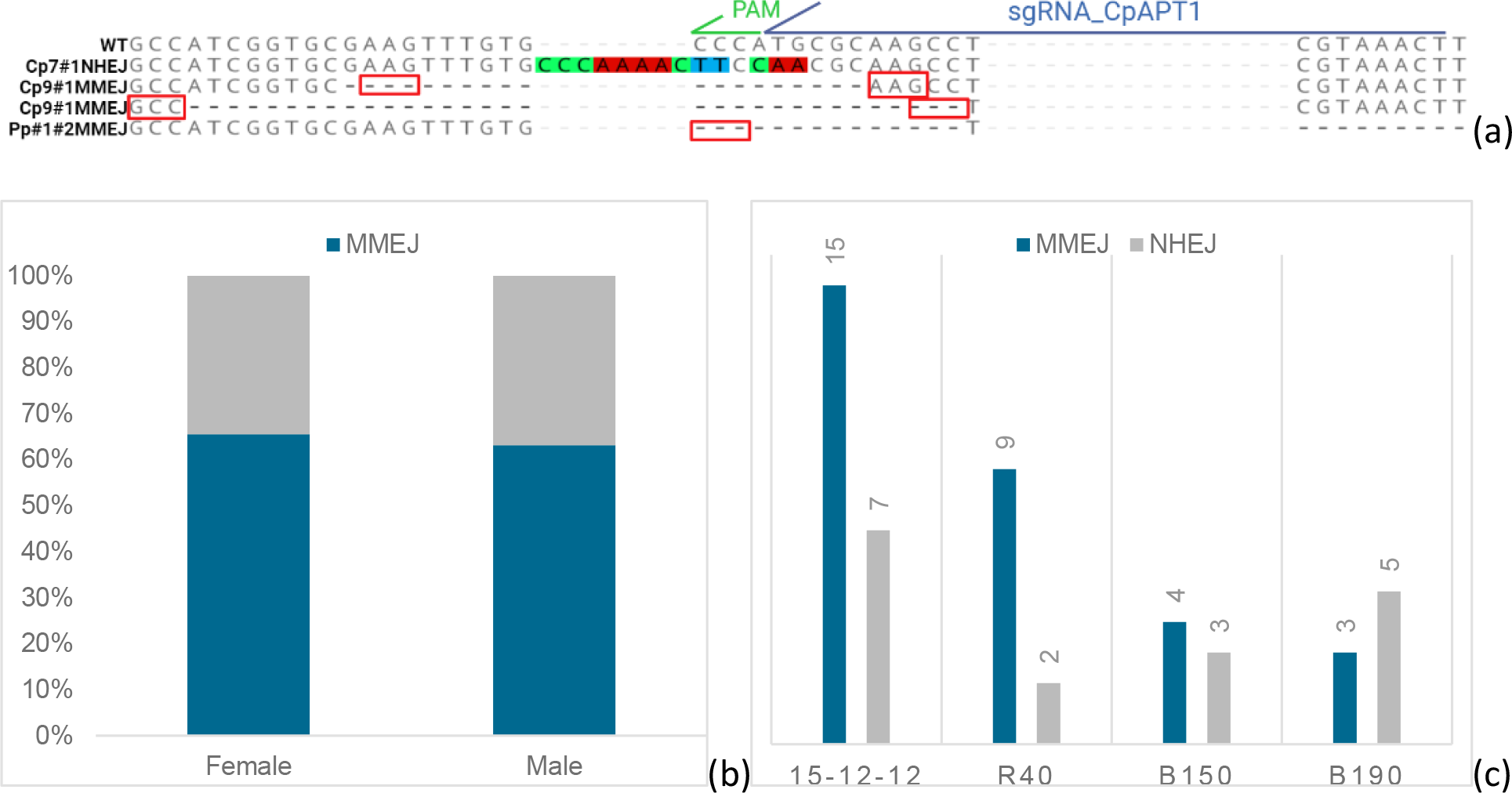
MMEJ and cNHEJ DSB repairs made in *Ceratodon purpureus*. a) Detailed examination of cNHEJ versus precise MMEJ repairs made at cut site CpAPT1 compared to the wildtype, including mutant Pp#1#2MMEJ which uses MMEJ to repair a double sgRNA deletion across *CpAPT*. b) Average percent MMEJ to NHEJ repairs in B150/15-12-12 (female) isolates and B190/R40 (male) isolates. c) Number of *C. purpureus* APT-KO mutants per moss isolate and type of DSB repair observed from sanger sequencing.

### *Similar to* P. patens, C. Purpureus *APT-KO plants display abortive gametophores*

Wildtype *C. purpureus* can survive 2-FA exposure to concentrations under 50µm. In comparison, *P. patens* wildtype can survive 2-FA concentrations under 10µm. We exposed *CpAPT* knock-out *C. purpureus* plants were resistant to 2-FA at concentrations of 50µM. Additionally, the plants showed no production of normal gametophores on agar. Mutant plants produced small, disorganized buds at the apex of rhizoids and did not generate mature gametophores. Neither the mutants where one nor both sgRNAs cut within CpAPT produced mature gametophores (Figure 5). All plants selected for phenotypic analyses can be found in Table S5.

**Figure 5.**
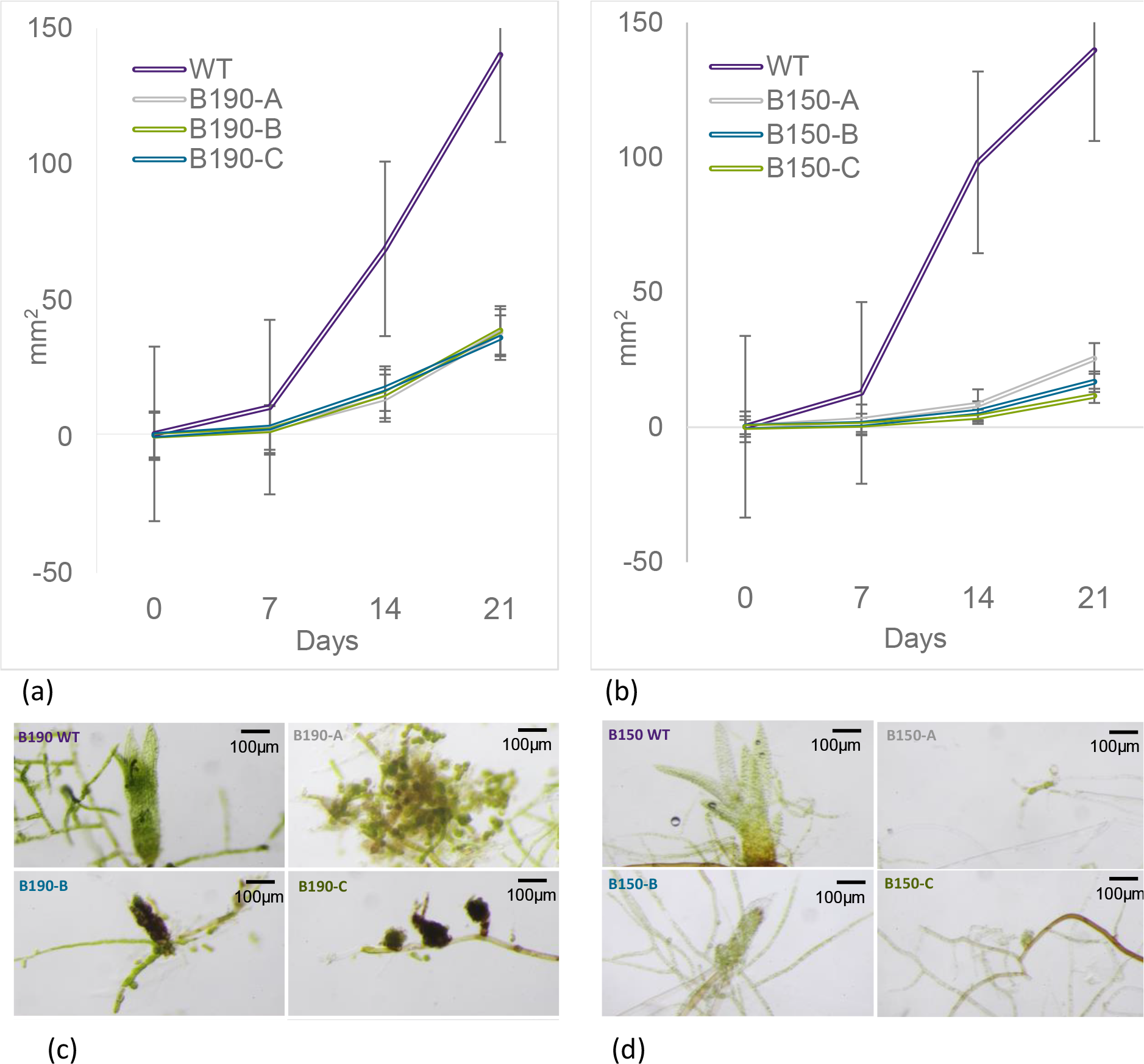
Area and stunted phenotype of APT-KO C. purpureus mutants. A) Three mutants and the wildtype male *C. purpureus* B190 average areas of growth over 21 days on PpNH_4_ media. B) Three mutants and the wildtype female *C. purpureus* B150 averages areas of growth over four weeks on PpNH_4_ media. C) B190 mutant and wildtype gametophore production after four weeks of growth. D) B150 mutant and wildtype gametophore production after four weeks of growth. Scalebar set to 100µm.

### *Efficient gene knock-in in* C. purpureus *with RNA-guided Cas9 nuclease*

To investigate whether CRISPR-Cas could be used to knock in transgenes in *C. purpureus*, we first chose a landing site in both male and female genomes. We selected 1000bp of gDNA that was in the vicinity, but not interrupting, two annotated genes (Figure S5). We verified the expression of the two flanking genes by using available jBrowse RNA-transcription data. The two landing pads, named CpLand1-GG1 and CpLand1-R40, can be found between 10716981-10717980 and 10985005-10986004 in the two respective R40 and GG1 genomes on chromosome 12. We designed sgRNA_CpLand1 to target the center of both landing sites.

For robust expression of two reporter genes, eGFP and mCherry, we looked for the equivalent of the well-established *A. thaliana* ribosomal protein of the 5a (RPS5a) subunit promoter in *C. purpureus* (Tsutsui and Higashiyama, 2017). We identified the *C. purpureus* RPS5a promoter using the *A. thaliana* RPS5a gene as a BLAST input. We selected the 1300bp upstream of the best hit result, CepurR40.3G229600, to be the promoter region. All promoters used in this study are available in Table S4 with promoter elements highlighted.

We co-transformed *C. purpureus* female (15-12-12) or male (R40) protoplasts by PEG-mediated transformation with pAct-Cas9, sgRNA_CpLand1, and either the eGFP or mCherry expression cassettes flanked by the 5’ and 3’ borders of CpLand1-GG1 or CpLand1-R40 (CpLandGFP-R40, CpLandGFP-GG1, CpLandmCherry-R40, CpLandmCherry-GG1). From the 50 potentially transformed 15-12-12 plants we screened per reaction, we confirmed one insert of GFP and mCherry respectively into the landsite, resulting in a 2% successful HDR rate. We verified both flanks of the recombined reporter genes were inserted correctly into the landsite, however, we were unable to amplify the entire gene from the 5’ to 3’ end (Table S2, Figure S4). In addition to PCR, we detected each constructs’ respective fluorescence with a Zeiss Axiozoom, and later confocal projections imaged on Zeiss LSM 710 confocal microscope (Figure 6). The selected endogenous CpRpS5a promoter does indeed express GFP and mCherry in *C. purpureus*.

**Figure 6.**
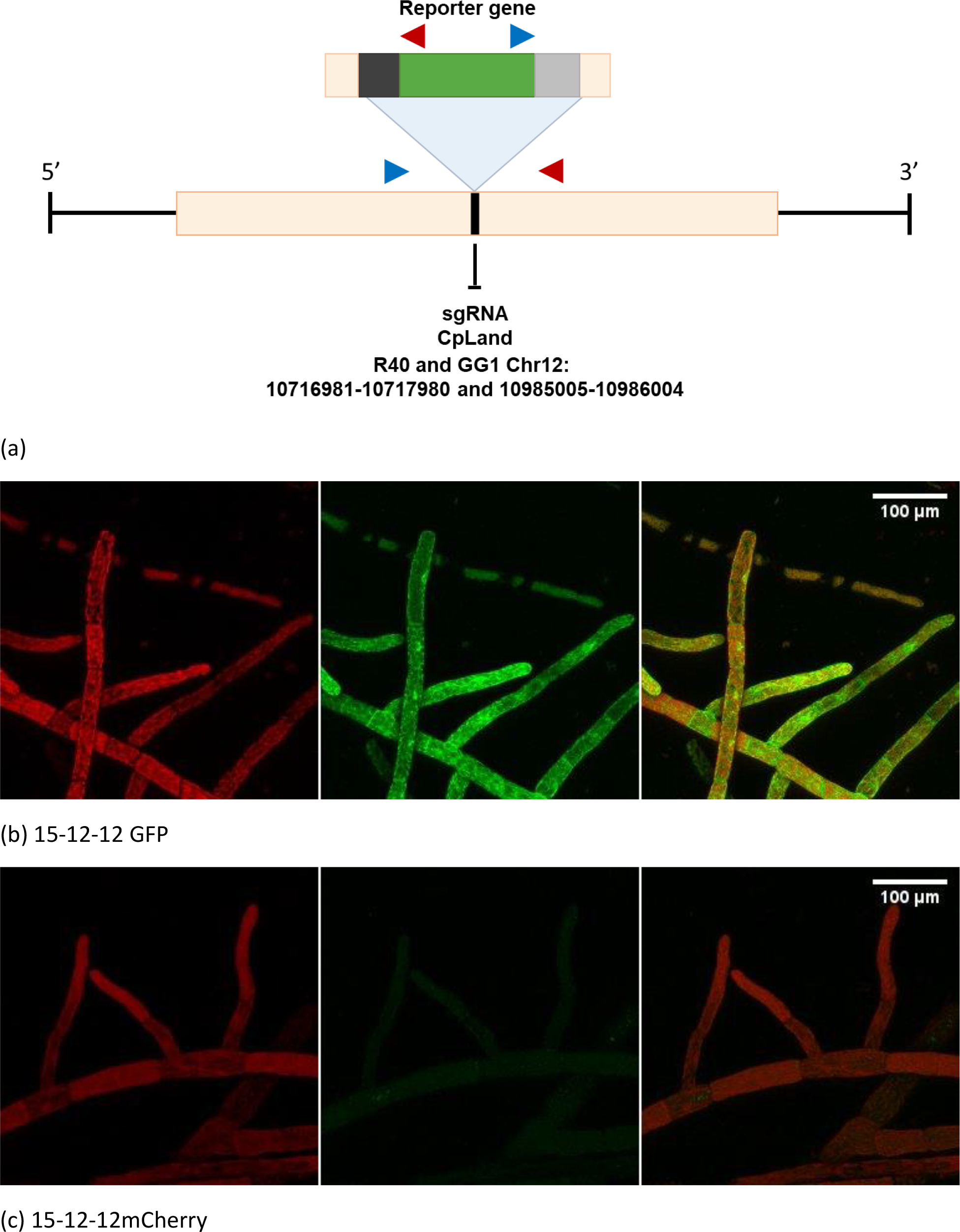
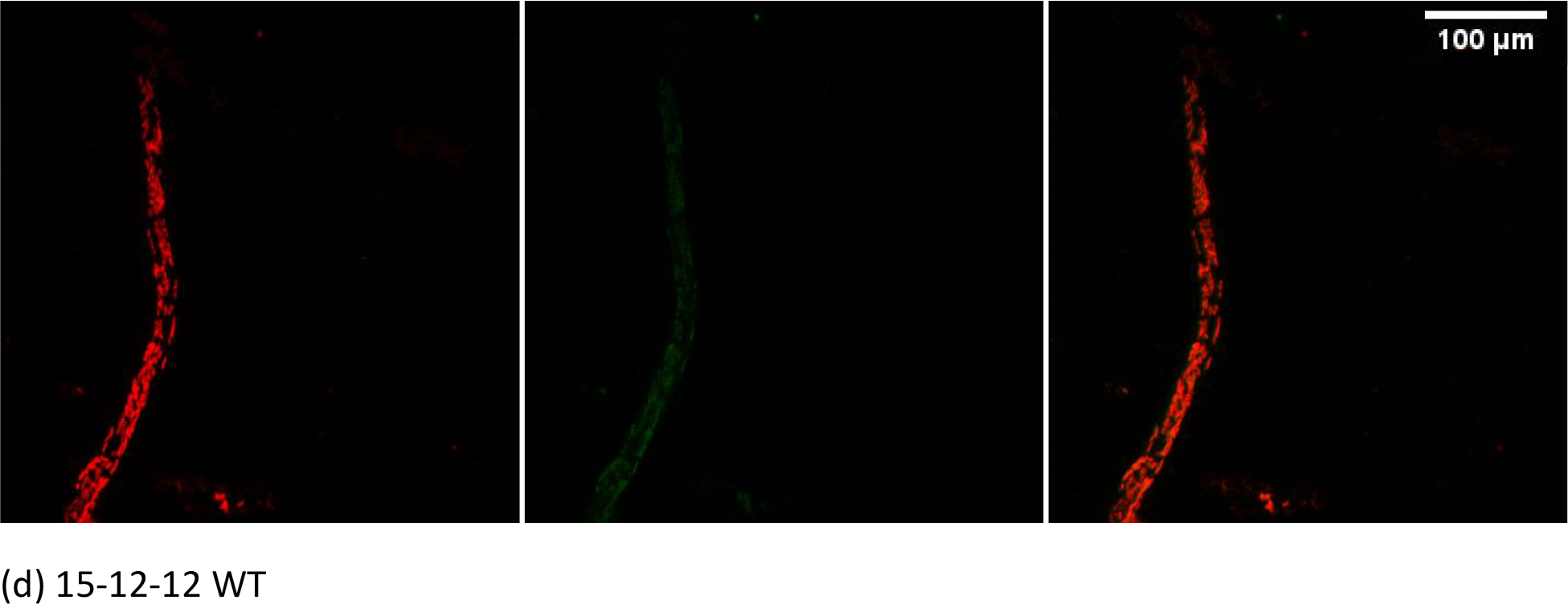
Scheme of CpLand and confocal maximum projections of C. purpureus 15-12-12 female isolates expressing mCherry and GFP in CpLand, compared to the wildtype. A) CpLand genomic location with sgRNA targets and primer annealing locations marked. 20bp sgRNA target site, CpLand, is indicated in black. Forward primers (blue) and reverse primers (red) show which regions were amplified with PCR and sequenced. B) a) 15-12-12 chloroplast autofluorescence, expression of GFP, and overlay of chloroplast and autofluorescence. C) Female 15-12-12, expression of mCherry, chloroplast autofluorescence, and overlay of chloroplast and autofluorescence. D) WT 15-12-12 chloroplast autofluorescence on the mCherry and GFP wavelengths, and an overlay of both autofluorescence wavelengths. The scalebar is set to 100µm in all confocal projections.

We selected and PCR screened 50 potential mutants per reaction of R40 male *C. purpureus* plants that we transformed with αCpLandR40-GFP or αCpLandR40-mCherry, with sgRNA_CpLand1 and pAct-Cas9. From these plants, we did not identify any transgenic plants correctly recombined with the reporter genes in the landsite.

## Discussion

Mosses are developmentally and biochemically highly diverse, but nonetheless remain a largely unexplored resource for novel gene analyses. Reverse genetic studies of gene function are well established in *P. patens*, but the potential for gene targeting in other mosses remains untapped. CRISPR-Cas technologies now enable transgenic research beyond traditional model systems. Here we used the previously established *APT* knock-out selection screen (Trouiller et al., 2007) to test the functionality of CRISPR-Cas9 gene editing components in the dioecious moss, *C. purpureus*. We found that three of the four native *C. purpureus* U6 promotors that we tested generated more edits than the foreign *P. patens* U6 promotor, but we also found substantial variation among sgRNA targets and among *C. purpureus* genotypes. Because the common ancestor of *C. purpureus* and *P. patens* is shared with ∼95% of all moss species (Bechteler et al., 2023; Laenen et al., 2014), the molecular tools we evaluated are likely to function across much of moss diversity. Additionally, we established a novel landsite for the ectopic expression of foreign genes inserted in the *C. purpureus* genome using HDR. Collectively these analyses demonstrate that CRISPR can be deployed for functional analyses across mosses, and to provide novel insights into land plant evolution (Bechteler et al., 2023).

### *Overcoming Barriers to Gene Knock Outs with CRISPR-Cas in* C. purpureus

The constructs transcribed under the U6 promoter Cp10 had the greatest GE frequency of edited plants (Figure 2). This increase in targeting efficiency is potentially due to its characteristic USE found -25bp upstream of the TATA box. An optimal U6 promoter maintains 25bp space between the TATA box and the USE to properly bind the RNA Pol III (Wang et al., 2008). While all the promoters we evaluated contained both regulatory elements, the spacing between these elements may explain the variation in efficacy (Ma et al., 2014;Table S4).

While the *P. patens* promoter also maintains a 25bp gap between the TATA box and USE transgene promoters from other species can be highly methylated compared to endogenous promoters. (Long et al., 2018; Riu et al., 2023; Wang et al., 2008). We tested four native *C. purpureus* U6 sgRNA promoters against one exogenous promoter from *P. patens* with sgRNA constructs targeting the *CpAPT* gene. This organismal preference for native promoters may explain the difference between efficiencies of editing of *Pp*U6 and *Cp10*U6.

Another explanation of the difference in editing efficiency between constructs is due to variable transfection efficiencies. To eliminate confounding factors, we isolated the protoplasts from tissues grown under the same conditions, all plasmids ranged from 5 – 9 kbp (Figure S1), we transformed the protoplasts with normalized plasmid concentrations and PEG molecule size, we heat shocked all cells at the same temperature and for the same duration, and we regenerated the transformed protoplasts under the same conditions.

We also found consistent differences in performance between guides across all isolates (Figure S4). We predicted that CpAPT2 would have a greater efficacy by U6 transcription predictions of Doench et al 2014 in CRISPOR with a score of 57 for CpAPT1 and 79 for CpAPT2. (Hassan et al., 2021; Doench et al., 2014; Waibel and Filipowicz, 1998). Both CpAPT1 and CpAPT2 use NGG protospacer adjacent motifs (PAM) specified for *Streptococcus pyogenes* Cas9 recognition, and there is no evidence of a favoring of the CGG over the TGG PAM for Type II Cas9 (Anders et al., 2014). Natural variation in sgRNA efficiency exists between all crRNAs due to epigenetic regulation or DNA structural differences (Wu et al., 2014). This variation may explain the evident difference in guide efficacy.

Remarkably, we classified 52% of the *C. purpureus* APT-KO plants as mosaics, containing multiple distinct edits. it is possible that the apparent mosaics represent the chance isolation of two edited protoplasts that grew together. However, we assured the initial density of protoplasts plated on the petri dishes was low, with ∼660-1200 protoplasts (each 30-35µm in diameter) per cm^2^, and we isolated transformants under a dissecting microscope. Alternatively, the mosaics may represent distinct cells in a single individual that were edited separately during early development. If true, Cas9 activity must have been initiated after the first cell division. This high frequency of mosaics suggests that *C. purpureus* may have a wide window during which cells can be edited by CRISPR-Cas9, and that the Cas9 may continue to be active after the initial cell division. Chimeric plants may be genetically separated by undergoing enzymatic digestion and regeneration a second time.

It is possible the rice actin promoter used to express the Cas9 could be less efficient in the *C. purpureus* genomic background compared to *P. patens*. We suggest future experimentation in *C. purpureus* to choose a tissue-specific promoter that is active in the early stages of protonemal development to express the Cas9. Early expression of the Cas endonuclease may help ensure the DSB is cleaved before the moss cell undergoes its first division.

In addition to variability in performance of the U6 promotors and sgRNAs, we observed variation among *C. purpureus* genotypes. In the *CpAPT* knock-out trials, the female isolate, 15-12-12, had a lower overall editing efficiency than the male isolate, R40 (Figure 3). It is important to note we tested only one female isolate and one male isolate of *C. purpureus* in the editing frequency and regeneration calculations. Interestingly, the R40 male genome has 36 copies of various RNA polymerase III subunits with six of these sequestered on the male V sex chromosome. In contrast, the GG1 female genome has 34 subunit orthologs, with two copies on the female U sex chromosome (Table S6). Comparably, *P. patens* maintains 32 RNA polymerase III subunit orthologs. We do not yet know if higher RNA polymerase III copy number translates to more sgRNA transcription and higher editing efficiency, but it is one explanation for the differences seen between R40 and 15-12-12. Given that the species is highly polymorphic, it is unlikely that the exact gene copy number will be consistent across the species. As we have seen even here, B190 was the only isolate to maintain a DSB and repair at CpAPT1 (Figure S4).

Of the plants that have been studied, most repair DSBs generated by environmental stressors or CRISPR-Cas targeting using the non-homologous end joining (cNHEJ) pathway (Pawelczak et al., 2018). In addition to cNHEJ, *P. patens* also employs microhomology-mediated Alternative EJ (Alt-EJ) (also known as MMEJ) and homology-directed repair (HDR) (Collonnier et al., 2017) to repair DSBs. *C. purpureus* repaired CRISPR-Cas DSBs with all three pathways and favored Alt-EJ for most repairs. The importance of Alt-EJ in repair of DSBs in *P. patens* and *C. purpureus*, suggests this repair pathway may have been inherited from the common ancestor of these mosses.

### *CRISPR-Cas can be used for the targeting of transgenes in* C. purpureus

A key tool for gene functional analysis is a landing site for the reliable ectopic expression of foreign genes. Other methods used to incorporate HR plasmids, such as particle bombardment and agrobacteria mediated transformation, may randomly integrate unwanted segments of the HR cassettes in the genome and may negatively disrupt coding regions (Altpeter et al., 2016). Moss protoplasts can be readily induced to undergo gene targeting using HDR by a direct PEG-mediated transformation protocol. Targeted gene knock outs using HDR has been used in *C. purpureus* (Trouiller *et al*., 2007), but we show here improvements in HDR efficiency of transgene integration events into a known location by combining HR plasmids with CRISPR-Cas9. We detected HDR-mediated gene editing by the expression of mCherry and GFP fluorescence in the selected landsite in *C. purpureus* isolate 15-12-12 at a rate of 2%. Previous GT events in *C. purpureus* using HDR in Trouiller et al., 2007 ranged from 0.5-1.5% without the use of CRISPR-Cas9. Overall, GT frequencies in the moss *C. purpureus* are still lower than the typically displayed efficiency of 20% for *P. patens*, but much higher than rates of 0.1% demonstrated in angiosperms (Sun et al., 2016; Svitashev et al., 2015).

HR gene insertion is still functional with genetic distance between the homology arms and the genome. We constructed the homology arms of the male isolate, R40, using the published R40 genome, and we confirmed with Sanger sequencing that there were no SNPs present between the plasmid homology arms and the genome. Conversely, we constructed the homology arms for the female isolate, 15-12-12, from the published female genome, GG1. There were many SNPS as well as insertions between the plasmid and the genome, however the constructs still successfully integrated the reporter genes (Figure S7), which were expressed by the *C. purpureus RPS5a* gene promoter. Not finding an R40 mutant expressing a reporter gene may be attributed to the small sample size of 50 mutants screened. Additionally, the use of alternative genomic locations in the future to compare landsite may increase HDR rates.

## Conclusion

Here we have vetted tools for CRISPR-Cas9 gene knock-outs and knock-ins in *C. purpureus*, including identifying an effective U6 promotor, showing that the *P. patens* U6 promotor works across 200 million years of evolution, established a new landing site for ectopic expression, and tested a new promotor for driving expression in juvenile and potentially mature tissues in this species. These methods established in this study pave the way for expanding techniques such as base and prime editing (Guyon-Debast et al., 2021; Perroud et al., 2022) in *C. purpureus*. While screening may be more time intensive due to fewer successful transformants in *C. purpureus* compared to *P. patens*, the regeneration time of moss is short (2-3 months from protoplast isolation to DNA extraction) and HDR mutants may be more easily identified than in angiosperms with frequencies lower than 1% (Sun et al., 2016; Svitashev et al., 2015). This work provides a promising future of studies to be done in *C. purpureus* genomic repair mechanisms and opens avenues for further analysis of sex-specific gene function and evolution in plants.

### Experimental Procedures

#### Bioinformatics, Molecular Cloning and Vectors

We used the https://phytozome-next.jgi.doe.gov/ Basic Local Alignment Search Tool (BLAST) and CRISPOR 5.01 (Concordet and Haeussler 2018) for gene identification and to select the best guide RNA to avoid off-target effects.

The pAct-Cas9 plasmid used in this study contains a Cas9 expression cassette under control of the rice actin 1 promoter and a codon-optimized version of Cas9 from *Streptococcus pyogenes* fused to a SV40 nuclear localization signal (Collonnier et al., 2017a).

For expression of the sgRNAs, we identified *C. purpureus* genomic sequences for the four U6 genes, CpU6-7 (coordinates 11559096-11558782 on chromosome 7), CpU6-9 (coordinates 3874021-3873707 on chromosome 9), CpU6-10 (coordinates 13131293-13130980 on chromosome 10 and CpU6-12 (coordinates 3954576-3954890 on chromosome 12) by Basic Local Alignment Search Tool (http://www.phytozome.net/physcomitrella_er.php) using the *A. thaliana* U6-26 snRNA sequence (X52528; Li et al., 2007) as the query.

We defined two 5’-N(20)-3’ sequences to target and knock out the *CpAPT* gene. We combined the synthesis of the 20bp target with the tracrRNA scaffold (Chen et al., 2013). Each sgRNA was expressed by one of the four different U6 promoters from *C. purpureus*, or the *P. patens* U6 promoter (Collonnier et al., 2017a). Each fragment containing the promoter, 20bp target, and tracrRNA was synthesized by TwistBioscience, San Francisco, CA, USA.

We introduced these different sgRNA cassettes into the pDONR207-neomycin resistant (NeoR) plasmid (Guyon-Debast et al., 2021) using Gateway™ BP Clonase II (ThermoFisher Scientific, USA) to get the plasmids sgRNA_CpAPT1_#1 to #5 and sgRNA_CpAPT2_#1 to #5.

To integrate genes into the landing sites, we synthesized GoldenBraid (GB) compatible α entry clones, CpLandR40 and CpLandGG1, containing 1000bp of homologous sequence specific to the R40 (Chr12:10716981-10717980) and GG1 (Chr12:10985005-10986004) *C. purpureus* genomes and cloned them into a pTwist Amp High Copy plasmid (TwistBioscience, San Francisco, CA, USA). Using GB cloning standard multipartite assembly, we constructed CpLandmCherry-R40, CpLandmCherry-GG1, CpLandGFP-R40, and CpLandGFP-GG1 using three parts: the promoter of the *C. purpureus* native RpS5a (CpurR40 Chr3:19311367-19312667), CDS of the selected reporter genes, eGFP (C5MKY7) or mCherry (X5DSL3) respectively, and a terminator sequence of the *Pisum sativum* ribulose bisphosphate carboxylase small subunit (RbcSE9) (Sarrion-Perdigones et al., 2011). To generate sgRNA_CpLand1, we designed the 5’-N(20)-3’ sequence to target both CpLand-R40 and CpLand-GG1 (Figure S6) landsites and used CRISPOR 5.01 to minimize offsite effects (Concordet and Haeussler 2018). We synthesized this 20bp target and tracrRNA scaffold (Chen et al., 2013) under expression of the *P. patens* U6 promoter (Collonnier et al., 2017a) (TwistBioscience, San Francisco, California, USA). We introduced this sgRNA cassette into NeoR (Guyon-Debast et al., 2021) using Gateway™ BP Clonase II (ThermoFisher Scientific, USA). All sgRNA sequences are available in Table S1. All maps of plasmids used are available in Figure S1.

#### Plant material and transformation

We cultivated and vegetatively propagated *C. purpureus* strains B150 and B190 (isolated from Yukon, Alaska), R40 (isolated from Rensselaer, NY, USA; McDaniel et al. 2013) and 15-12-12 (isolated from Storrs, Connecticut, USA) according to Cove et al., 2009; Schaefer and Zrÿd, 1997. We grew the cultures at 60% humidity, 25°C with a 16 h light (70–80 µmol/m^2^/s)/8 h dark cycle.

We isolated moss protoplasts and transformed them as previously described (Charlot et al., 2022) with the following modifications. We propagated vegetative protonemal tissues on PpNH_4_ media with overlain cellophanes for 4-5 days. We then enzymatically digested tissues for 45 minutes in low light using 0.5% Driselase (Sigma Aldrich, D9515) with occasional agitation. We filtered the protoplast suspension using 32-40µm steel mesh filters, and rinsed protoplasts at all rinse steps with a solution of 8.5% mannitol-7mM CaCl_2_.

To knock out the *APT* gene, we co-transformed an estimated 1.5 to 2 x 10^5^ protoplasts in MMM buffer with 6µg of pAct-Cas9 (Collonnier et al., 2017a), 6µg sgRNA_CpAPT1_#1, 6µg and sgRNA_CpAPT2_#1 plasmids at 1000ng/µl each. We repeated this for sgRNA-CpAT1_#2 to #5, and sgRNA_CpAPT2_#2 to #5. The transformed protoplasts are mixed 1:1 with a 1.5% alginic acid (Sigma Aldrich, USA), 8.5% mannitol, 7mM CaCl_2_ solution and plated on PpNH_4_ solid medium supplemented with 8.5% mannitol overlaid with sterile cellophane. The plates are left at 25°C in the dark for 24 hours before being moved to a 25°C long-day, light cycle growth chamber. After 5 days of growth, we transferred the cellophane disks to PpNH_4_ containing 50µM final concentration of the toxic adenine analogue, 2-fluoroadenine (2-FA, Fluorochem, UK) suspended in DMSO. We counted and isolated the surviving plants after 12 days of exposure to 2-FA. We picked the surviving plants and transferred them to solid PpNH_4_ medium to grow for 10 days before harvesting them for DNA extraction.

To insert the reporter genes by HDR into the CpLand-R40 and CpLand-GG1 sites, we co-transformed an estimated 1.5 to 2 x 10^5^ protoplasts in MMM buffer with 6µg of pAct-Cas9, 6µg of sgRNA_CpLand1, and 6µg of CpLandmCherry-R40, CpLandmCherry-GG1, CpLandGFP-R40, or CpLandGFP-GG1 respectively. The transformed protoplasts are mixed 1:1 with a 1.5% alginic acid (Sigma Aldrich, USA), 8.5% mannitol, 7mM CaCl_2_ solution and plated on PpNH_4_ solid medium supplemented with 8.5% mannitol overlaid with sterile cellophane. The plates are left at 25°C in the dark for 24 hours before being moved to a 25°C long-day, light cycle growth chamber. After 5 days of growth, we transferred the cellophane disks to PpNH_4_ containing a final concentration of 25 mg/l G418 antibiotic (Duchefa, Netherlands). We isolated surviving plants after 12 days of exposure to G418 and transferred to PpNH_4_ medium to grow for 10 days before being harvested for DNA extraction.

#### DNA extraction, PCR and Sanger Sequencing of mutant plants

We extracted genomic DNA from ∼50mg of fresh tissue from mutants and wild-type strains using SDS lysis buffer extraction methods described in Lopez-Obando et al., 2016. We resuspended DNA in 50-80µl of 8.0pH TE buffer and stored them at –20°C. We Sanger sequenced the candidate *CpAPT*-KO mutants (Genoscreen, France and Eurofins Genomics, USA) using primers CpAPTKO-1 and CpAPTKO-2 to amplify *CpAPT*. Similarly, we sequenced the CpLand1 inserts using the primers CpLand1-For and CpLand2-Rev. To confirm the insert was correct, CpLand1-For paired with CpLand-RpS5a-Rev verified the 5’ region of insertion, while CpLand-RbcSE9-For and CpLand2-Rev verified the 3’ region of insertion (Figure 1). All primers were designed using NCBI Primer BLAST (https://www.ncbi.nlm.nih.gov/tools/primer-blast/; Ye et al. 2012) and are available in Table S2. All Sanger sequencing data is available in Data S1.

#### Imaging and phenotype evaluation

We imaged the B150 and B190 *C. purpureus* WT isolates and three different APT-KO mutants of both isolates at days 0, 7, 14, 21 at 10x and 7x magnifications with a Canon EOS 50D DSLR camera and Olympus SZX16 stereomicroscope setup. After 21 days, the plants became too large to encompass in the lowest magnification setting of 7x. We grew each plant starting from ∼4-12 cells on 20ml of PpNH_4_ solid agar media in 15mm deep petri dishes. To account for possible edge effects, we observed mutants grown alongside their wild type counterparts on two different layouts (Figure S2). We analyzed all photos using ImageJ to calculate the area of each isolate, and used two separate macro scripts to analyze photos depending on if the plant was imaged using bright field or dark field microscopy filters. All macro scripts can be found in Table S3. We screened the R40 and 15-12-12 *C. purpureus* CpLand1 transgenic candidates with fluorescence microscopy using a Zeiss Axio Zoom.V16 Stereo Microscope. We imaged the Sanger sequenced 15-12-12 expressing the correct fluorescent inserts using a Leica Sp5 confocal laser fluorescent microscope.

### Accession numbers

## Acknowledgments

The authors would like to express their sincere gratitude to all individuals who have contributed to the successful completion of this research. Special thanks to Julie Calbry, Florence Charlot, and Anouchka Guyon from the INRAE, IJPB in Versailles for their valuable contributions in data collection. We would also like to acknowledge Mike Gunter, Pete Ryschkewitsch, and Jay Wheeler and Hua Yan of the Biology Department at the University of Florida for their continued assistance and support of this research, and for their expertise and guidance. Finally, we are grateful to the funding agencies, the Evolution Society, the Biology Department of UF, the University of Florida International Center, the IJPB, and the NSF for their financial support, which enabled the successful execution of this project. The IJPB benefits from the support of Saclay Plant Sciences-SPS (ANR-17-EUR-0007).

## Notes

https://zenodo.org/records/7979424

### Competing Interest Statement

The authors have declared no competing interest.

