## Supplemental Data for "Establishing CRISPR-Cas9 in the sexually dimorphic moss, *Ceratodon purpureus*"

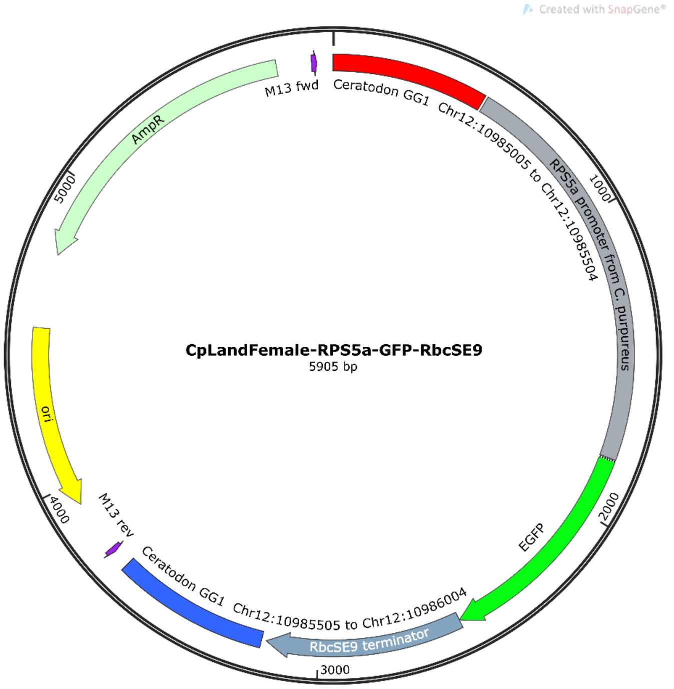


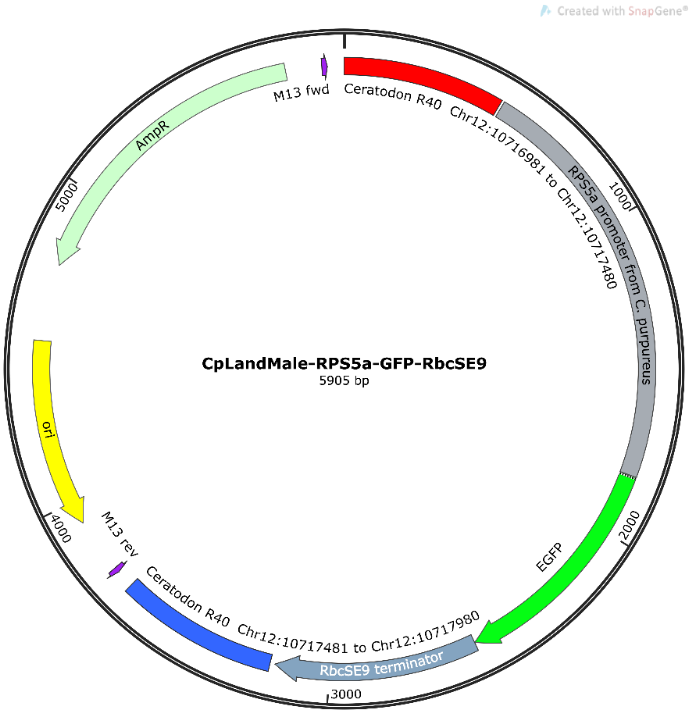

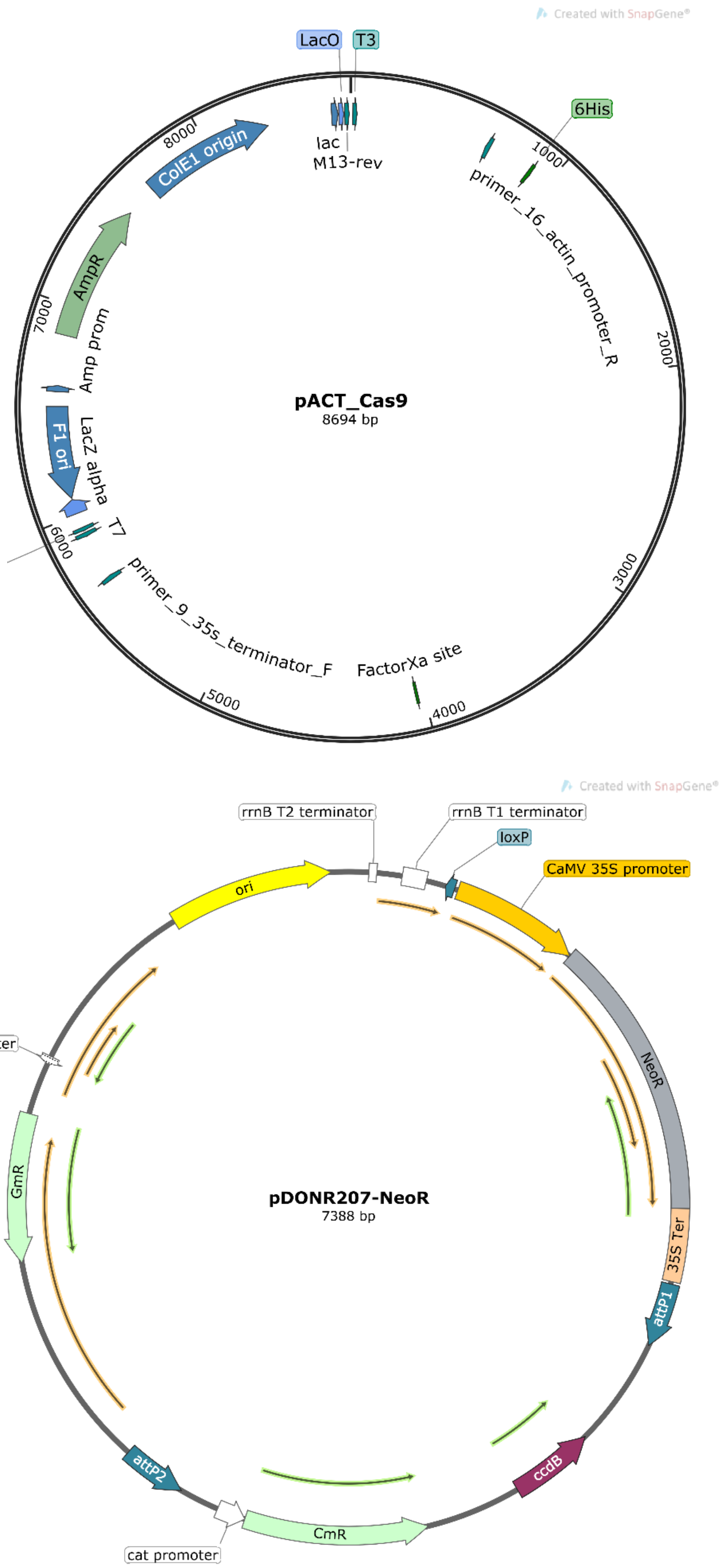


Figure S1. All plasmid maps of vectors which were used within this study.

**Layout 1.**

| | | | |
|---|---|---|---|
| B150/B190 WT | B150-A / B190-B | B150-B / B190-C | B150-C / B190 A |
| B150-C / B190 A | B150/B190 WT | B150-A / B190-B | B150-B / B190-C |
| B150-B / B190-C | B150-C / B190 A | B150/B190 WT | B150-A / B190-B |
| B150-A / B190-B | B150-B / B190-C | B150-C / B190 A | B150/B190 WT |

**Layout 2.**

| | | | |
|---|---|---|---|
| B150-B / B190-C | B150-A / B190-B | B150/B190 WT | B150-C / B190 A |
| B150-C / B190 A | B150-B / B190-C | B150-A / B190-B | B150/B190 WT |
| B150/B190 WT | B150-C / B190 A | B150-B / B190-C | B150-A / B190-B |
| B150-A / B190-B | B150/B190 WT | B150-C / B190 A | B150-B / B190-C |

Figure S2. Layout growth trials for *C. purpureus* APT-KO mutants and wildtype isolates: B150 (female) and B190 (male).

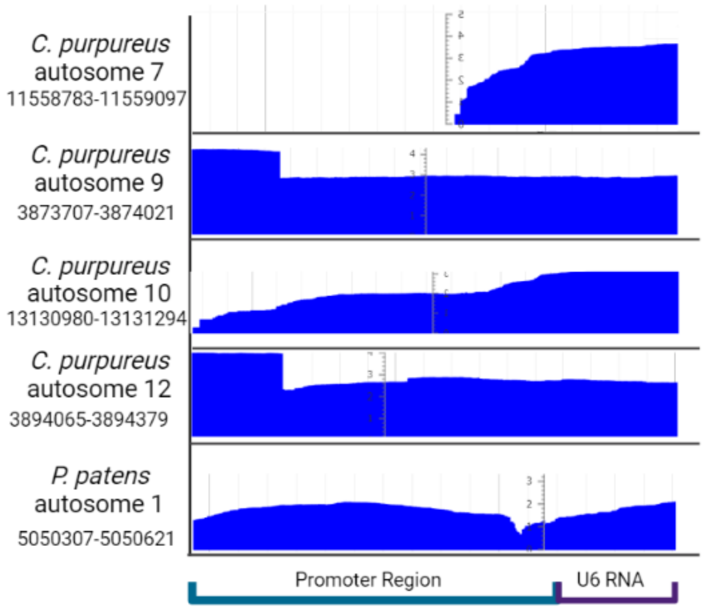


Figure S3. Native transcription of U6 snRNA in *C. purpureus* or *P. patens* from each U6 promoter used in this study.

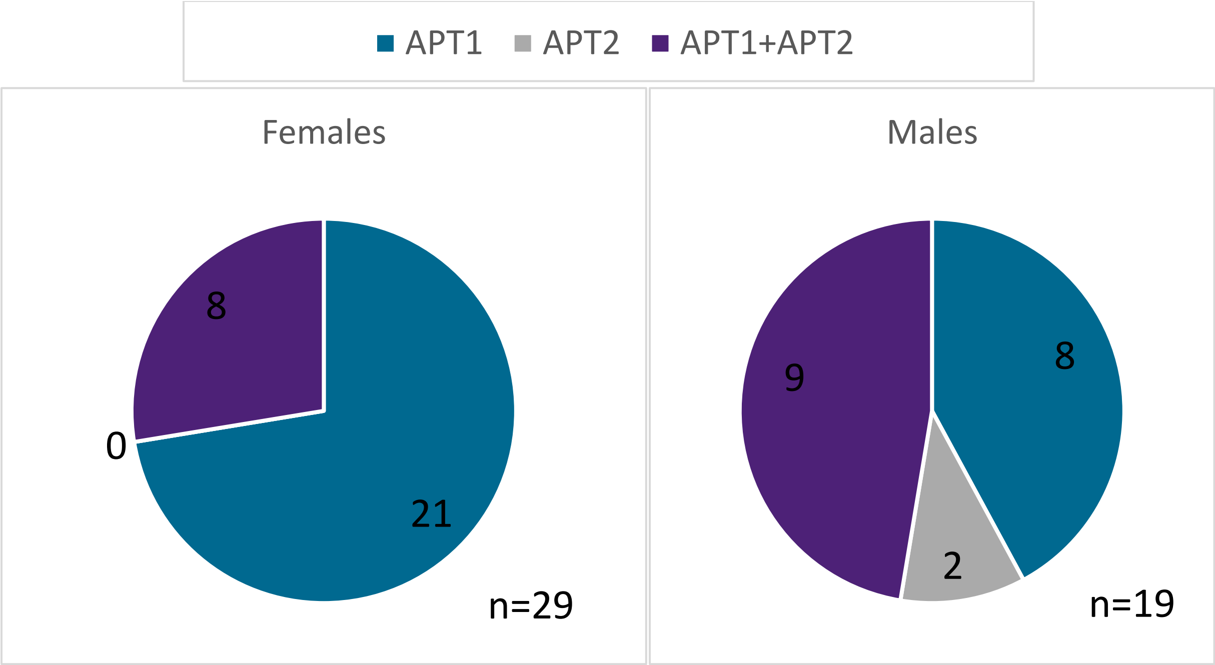


Figure S4. Number of cuts made by crRNA targets CpAPT1 and CpAPT2 in isolates B150/GG1 (females) and B190/R40 (males).

(a)

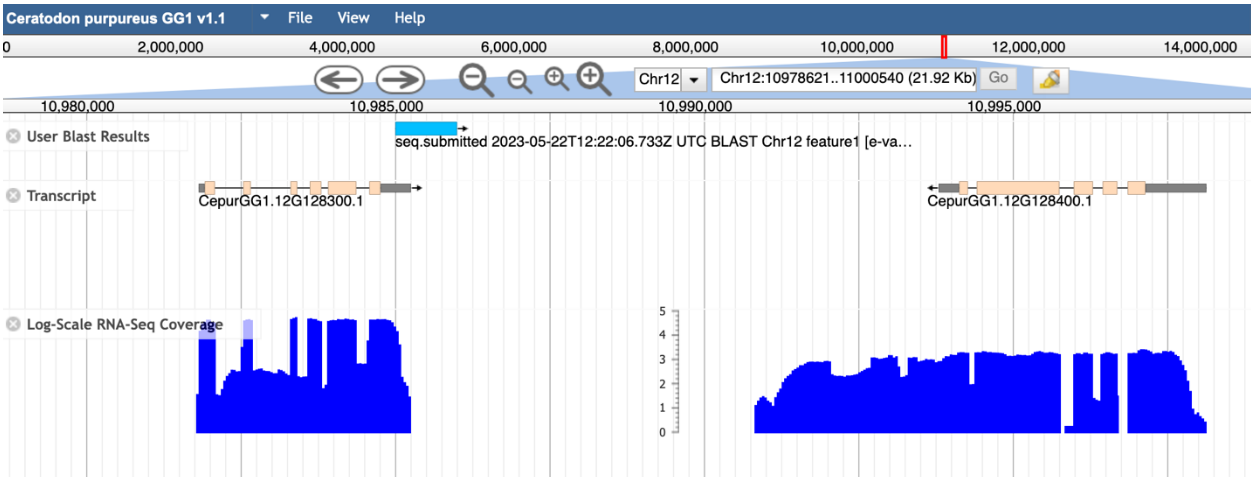


(b)

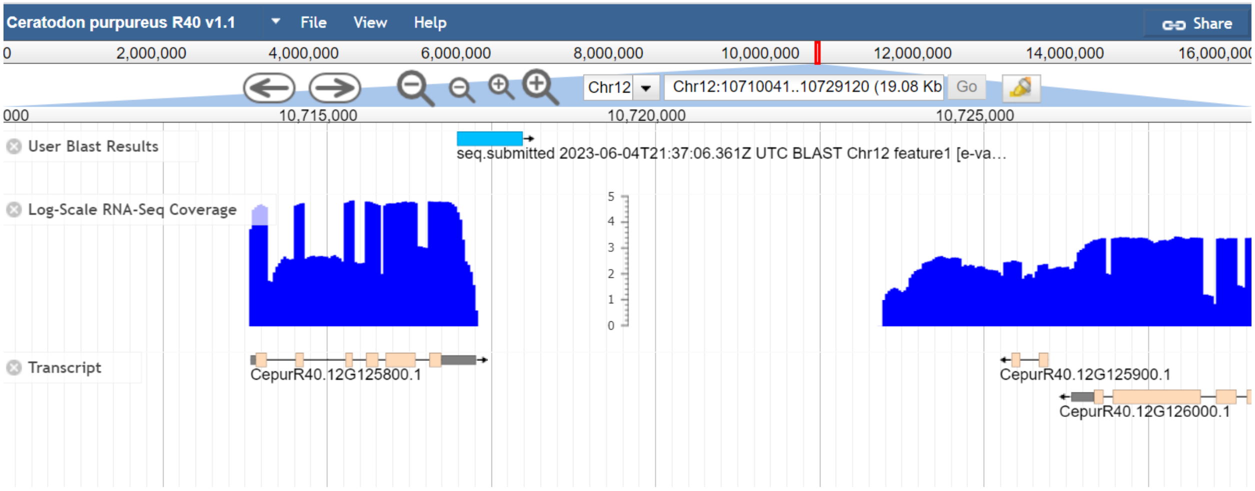


Figure S5. CpLand site shown to not be introduced within a coding region of either the (a) GG1 (female) or (b) R40 (male) genomic background.

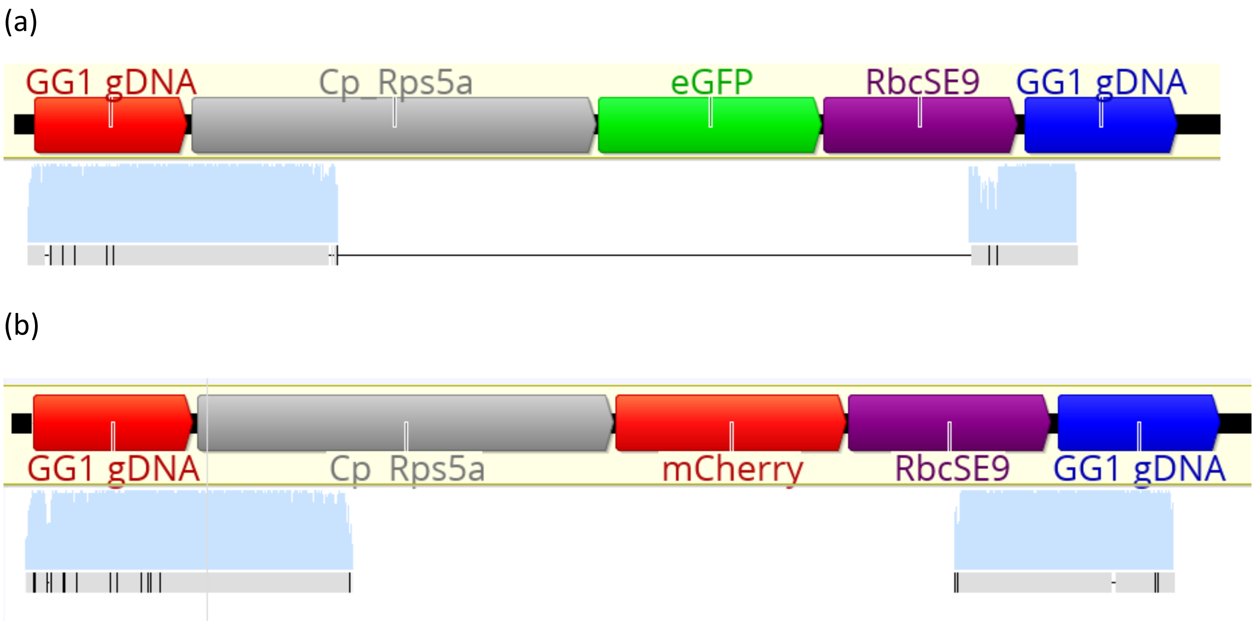


Figure S6. Alignment of CpLand inserts in *C. purpureus* 15-12-12 to the *C. purpureus* GG1 homologous arms and CpLandeGFP1 and CpLandmCherry1 plasmids. (a) eGFP insert with homologously recombined genomic flanks. (b) mCherry insert with homologously recombined genomic flanks.

(a)

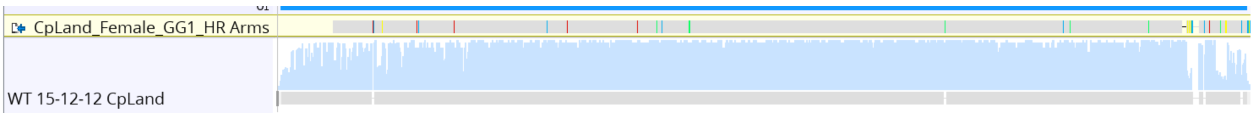


(b)

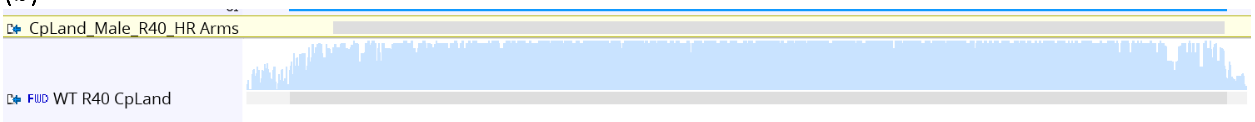


Figure S7. Wildtype alignments of CpLand site in 15-12-12 and R40 to the GG1 (a) and R40 (b) published genomes, respectively.

Table S1. All sgRNAs sequences that were used in this study with attB Gateway cloning sites in blue, U6 promoter sequence in red, crRNA 20bp sequence in black, the tracrRNA in yellow.

| |
|---|
| sgRNA-CpLand<br>*Physcomitrium patens* U6 promoter<br>ggggacaagtttgtacaaaaaagcaggcttcGTCCATTGAAGCAGACGTGTTGCGACAGGTTAGCGACGATGG<br>GTGTAGATGTGATGTGATGTGATGGTGTGGTTCTTCCACGGCGGCGTCCTTGCGGTGGCGGAGAAG<br>GGGATATCCCGAAGGAGCGGCAGCGGGAGAGCACAAGCAGAAAGGGTGCAGTGAGTGAGTGGGTC<br>CAGCTGGGTGGCTGGCCGAGTGGACGCGACCGGGTTTCGAGGGGGcGGGGGAGAAAAGGGATGGA<br>GCGAGGGATATAACCCACATGGAATGGAGGTGGGTGTGAAGGCGGGTATATAGGAAGGTGGAGGA<br>CTTACAACCCATgTTGCTACAGATGACTCAAGAGTTTAAGAGCTATGCTGGAAACAGCATAGCAAGTT<br>TAAATAAGGCTAGTCCGTTATCAACTTGAAAAAGTGGCACCGAGTCGGTGCTTTTTTTGTTTTTTATGT<br>CTgacccagctttcttgtacaaagtggtcccc |
| sgRNA_CpAPT1_#1<br>*Ceratodon purpureus* U6 promoter from autosome 7<br>ggggacaagtttgtacaaaaaagcaggcttcACCAAATCCTAAACGATAGTGAAACCAAATTTAGACGAAACAT<br>ATTCTCCAGCAGGTCATATTCTACTAGTAGTAATCTTCGCAGTGTAAACTAATGCTCTCATCTCCTAAA<br>AACGAGAAAAAAAGTAGTAAACTAAAGAAAGGAAGATGAGAAAGGACATTCTTCAGCAAAAGGGAA<br>CACAACTTCAACGCAAAACATGAGGAAAAAGCAAAAGTATTCTGACACAACACCTCTATTTTAGGCTG<br>ACAAAAACCATAACGTTTGTCACGACGTATACCGGCATCCATATAACCACAGACGGAACCTAACCTAT<br>TAGTTTACGAGGCTTGCGCATGTTTAAGAGCTATGCTGGAAACAGCATAGCAAGTTTAAATAAGGCTA<br>GTCCGTTATCAACTTGAAAAAGTGGCACCGAGTCGGTGCTTTTTTTGTTTTTTATGTCTgacccagctttctt<br>gtacaaagtggtcccc |
| sgRNA_CpAPT1_#2<br>*Ceratodon purpureus* U6 promoter from autosome 9<br>ggggacaagtttgtacaaaaaagcaggcttcATCTATGTCAAGGATGGACCACTCCTTTCCTTTCGCATCCACGA<br>GGAGTTGCAAGTGACGTGCAACCAAAGCGCCATCTCAAGGTGAGAAATTCGAATGTGGTCACTGCTG<br>CAAGTTGATTTTGCTCCCATTTTGACATACACATGTTTCCTTCACTGCAAGGAGTCTAGTAGGAATTTA |

| |
|---|
| TGTTGCCATACTTTAGAATCATAGGACTATTTCGCAGCAGTACAGTATCAGGGATGCACTTGTGATCT<br>GAAATCCCATAAGGATTGTGACTATAAACATAGCGAACTACTTAAGCTTGTGGTATCCTGCTTCCTTAT<br>AGTTTACGAGGCTTGCGCATGTTTAAGAGCTATGCTGGAAACAGCATAGCAAGTTTAAATAAGGCTA<br>GTCCGTTATCAACTTGAAAAAGTGGCACCGAGTCGGTGCTTTTTTTGTTTTTTATGTCTgacccagctttctt<br>gtacaaagtggtcccc |
| sgRNA_CpAPT1_#3<br>*Ceratodon purpureus* U6 promoter from autosome 10<br>ggggacaagtttgtacaaaaaagcaggcttcTTGCCATGACGAACTGTGGAATGAACCACCTAAGAGCTACTAT<br>CCTTGTTTATTTCCACTCGACTTATGTCATCTTGGAGTGGAAGTTAGAAGTAGTAGTTGGGTAGGCGA<br>CTCACTGACCCAGAGGATACCTTACGCAAGGAAAGGGACCCTGCGTGGGCAAGTTCACATTTGCCCT<br>GATGAAAAGCACACATACGGAGCATTGGGAAGGGCTGACATGGCATAGCATGGCATAGCATGGGAA<br>GTATAACCCATATCGTTTGGGCAGGGCAGAGAGGCGACCTATATAGTGCCGAGGGCGGGCTCTTCCT<br>TTCAGTTTACGAGGCTTGCGCATGTTTAAGAGCTATGCTGGAAACAGCATAGCAAGTTTAAATAAGGC<br>TAGTCCGTTATCAACTTGAAAAAGTGGCACCGAGTCGGTGCTTTTTTTGTTTTTTATGTCTgacccagcttt<br>cttgtacaaagtggtcccc |
| sgRNA_CpAPT1_#4<br>*Ceratodon purpureus* U6 promoter from autosome 12<br>ggggacaagtttgtacaaaaaagcaggcttcGCGAATGCCCAGAAGAAAGATGGCGGTTGTCGCACCTACATCA<br>TCGATGACAGTGATGCAGTGGGTGGTTGGGAAAGTGATGAGGTATGCCAAGACCTGCATACTGAGT<br>CAAACGACAGCCATGTCTGTTTTAAAGTTTTGCTGTTCTACCGGTACTCAAATCCCAGAAATGATAAGT<br>TGTTTCTTTTTCTGCAAGGTGCTGGAGTGGCATTAACTAAGATGGATACAGAGGATCGCATAATTGTC<br>ATATATACCCTAGGGTTAGCAACACTTTCGGAACCGTAATATATATTACCCCAAACCCCTCATCCTTTC<br>AGTTTACGAGGCTTGCGCATGTTTAAGAGCTATGCTGGAAACAGCATAGCAAGTTTAAATAAGGCTA<br>GTCCGTTATCAACTTGAAAAAGTGGCACCGAGTCGGTGCTTTTTTTGTTTTTTATGTCTgacccagctttctt<br>gtacaaagtggtcccc |
| sgRNA_CpAPT1_#5<br>*Physcomitrium patens* U6 promoter<br>ggggacaagtttgtacaaaaaagcaggcttcGTCCATTGAAGCAGACGTGTTGCGACAGGTTAGCGACGATGG<br>GTGTAGATGTGATGTGATGTGATGGTGTGGTTCTTCCACGGCGGCGTCCTTGCGGTGGCGGAGAAG<br>GGGATATCCCGAAGGAGCGGCAGCGGGAGAGCACAAGCAGAAAGGGTGCAGTGAGTGAGTGGGTC<br>CAGCTGGGTGGCTGGCCGAGTGGACGCGACCGGGTTTCGAGGGGGcGGGGGAGAAAGGGATGGA<br>GCGAGGGATATAACCCACATGGAATGGAGGTGGGTGTGAAGGCGGGTATATAGGAAGGTGGAGGA<br>CTTACAACCCATAGTTTACGAGGCTTGCGCATGTTTAAGAGCTATGCTGGAAACAGCATAGCAAGTTT<br>AAATAAGGCTAGTCCGTTATCAACTTGAAAAAGTGGCACCGAGTCGGTGCTTTTTTTGTTTTTTATGTC<br>Tgacccagctttcttgtacaaagtggtcccc |
| sgRNA_CpAPT2_#1<br>*Ceratodon purpureus* U6 promoter from autosome 7<br>ggggacaagtttgtacaaaaaagcaggcttcACCAAATCCTAAACGATAGTGAAACCAAATTTAGACGAAACAT<br>ATTCTCCAGCAGGTCATATTCTACTAGTAGTAATCTTCGCAGTGTAAACTAATGCTCTCATCTCCTAAA<br>AACGAGAAAAAAAGTAGTAAACTAAAGAAAGGAAGATGAGAAAGGACATTCTTCAGCAAAAGGGAA |

CACAACTTCAACGCAAAACATGAGGAAAAAGCAAAAGTATTCTGACACAACACCTCTATTTTAGGCTG
ACAAAAACCATAACGTTTGTCACGACGTATACCGGCATCCATATAACCACAGACGGAACCTAACCTAT
TGATTGCATCGAGATGCATGTGTTTAAGAGCTATGCTGGAAACAGCATAGCAAGTTTAAATAAGGCT
AGTCCGTTATCAACTTGAAAAAGTGGCACCGAGTCGGTGCTTTTTTTGTTTTTTATGTCTgacccagctttct
tgtacaaagtggtcccc

sgRNA_CpAPT2_#2
*Ceratodon purpureus* U6 promoter from autosome 9
ggggacaagtttgtacaaaaaagcaggcttcATCTATGTCAAGGATGGACCACTCCTTTCCTTTCGCATCCACGA
GGAGTTGCAAGTGACGTGCAACCAAAGCGCCATCTCAAGGTGAGAAATTCGAATGTGGTCACTGCTG
CAAGTTGATTTTGCTCCCATTTTGACATACACATGTTTCCTTCACTGCAAGGAGTCTAGTAGGAATTTA
TGTTGCCATACTTTAGAATCATAGGACTATTTCGCAGCAGTACAGTATCAGGGATGCACTTGTGATCT
GAAATCCCATAAGGATTGTGACTATAAACATAGCGAACTACTTAAGCTTGTGGTATCCTGCTTCCTTAT
GATTGCATCGAGATGCATGTGTTTAAGAGCTATGCTGGAAACAGCATAGCAAGTTTAAATAAGGCTA
GTCCGTTATCAACTTGAAAAAGTGGCACCGAGTCGGTGCTTTTTTTGTTTTTTATGTCTgacccagctttctt
gtacaaagtggtcccc

sgRNA_CpAPT2_#3
*Ceratodon purpureus* U6 promoter from autosome 10
ggggacaagtttgtacaaaaaagcaggcttcTTGCCATGACGAACTGTGGAATGAACCACCTAAGAGCTACTAT
CCTTGTTTATTTCCACTCGACTTATGTCATCTTGGAGTGGAAGTTAGAAGTAGTAGTTGGGTAGGCGA
CTCACTGACCCAGAGGATACCTTACGCAAGGAAAGGGACCCTGCGTGGGCAAGTTCACATTTGCCCT
GATGAAAAGCACACATACGGAGCATTGGGAAGGGCTGACATGGCATAGCATGGCATAGCATGGGAA
GTATAACCCATATCGTTTGGGCAGGGCAGAGAGGCGACCTATATAGTGCCGAGGGCGGGCTCTTCCT
TTCGATTGCATCGAGATGCATGTGTTTAAGAGCTATGCTGGAAACAGCATAGCAAGTTTAAATAAGGC
TAGTCCGTTATCAACTTGAAAAAGTGGCACCGAGTCGGTGCTTTTTTTGTTTTTTATGTCTgacccagcttt
cttgtacaaagtggtcccc

sgRNA_CpAPT2_#4
*Ceratodon purpureus* U6 promoter from autosome 12
ggggacaagtttgtacaaaaaagcaggcttcGCGAATGCCCAGAAGAAAGATGGCGGTTGTCGCACCTACATCA
TCGATGACAGTGATGCAGTGGGTGGTTGGGAAAGTGATGAGGTATGCCAAGACCTGCATACTGAGT
CAAACGACAGCCATGTCTGTTTTAAAGTTTTGCTGTTCTACCGGTACTCAAATCCCAGAAATGATAAGT
TGTTTCTTTTTCTGCAAGGTGCTGGAGTGGCATTAACTAAGATGGATACAGAGGATCGCATAATTGTC
ATATATACCCTAGGGTTAGCAACACTTTCGGAACCGTAATATATATTACCCCAAACCCCTCATCCTTTC
GATTGCATCGAGATGCATGTGTTTAAGAGCTATGCTGGAAACAGCATAGCAAGTTTAAATAAGGCTA
GTCCGTTATCAACTTGAAAAAGTGGCACCGAGTCGGTGCTTTTTTTGTTTTTTATGTCTgacccagctttctt
gtacaaagtggtcccc

sgRNA_CpAPT2_#5
*Physcomitrium patens* U6 promoter
ggggacaagtttgtacaaaaaagcaggcttcGTCCATTGAAGCAGACGTGTTGCGACAGGTTAGCGACGATGG

GTGTAGATGTGATGTGATGTGATGGTGTGGTTCTTCCACGGCGGCGTCCTTGCGGTGGCGGAGAAG
GGGATATCCCGAAGGAGCGGCAGCGGGAGAGCACAAGCAGAAAGGGTGCAGTGAGTGAGTGGGTC
CAGCTGGGTGGCTGGCCGAGTGGACGCGACCGGGTTTCGAGGGGGcGGGGGAGAAAAGGGATGGA
GCGAGGGATATAACCCACATGGAATGGAGGTGGGTGTGAAGGCGGGTATATAGGAAGGTGGAGGA
CTTACAACCCATGATTGCATCGAGATGCATGTGTTTAAGAGCTATGCTGGAAACAGCATAGCAAGTTT
AAATAAGGCTAGTCCGTTATCAACTTGAAAAAGTGGCACCGAGTCGGTGCTTTTTTTGTTTTTTATGTC
Tgacccagctttcttgtacaaagtggtcccc

Table S2. All primers used in this study and are written in the 5’ to 3’ orientation.

| | |
|---|---|
| CpAPTKO-1 For | TATGTTTCGGGATGTGACGA |
| CpAPTKO-2 Rev | CAACTTGGTCTGCATGATGG |
| CpLand-1 For | GTAACTCCATGGCGGGACAA |
| CpLand-2 Rev | TGGTTTATGCCAATGCTTGCT |
| CpLand-RpS5a-Rev | ATTTGGCCGGGTACGTTGAT |
| CpLand-RbcSE9-For | TGCTTTCGTTCGTATCATCG |
| CpLandBack_For | CCGCTTACCGGATACCTGTC |
| CpLandBack_Rev | GATTTGAAGCCCCCTACCCA |

Table S3. Image J macro scripts that were used to analyze the area of each moss image, whether imaged on either bright field or dark field.

| **Image taken on Bright-Field** | **Image taken on Dark-Field** |
|---|---|
| run("Convert to Mask"); | run("Select All");<br>run("Enhance Contrast...", "saturated=0.35");<br>doWand(3030, 288);<br>run("Subtract Background...", "rolling=50 light");<br>run("Subtract Background...", "rolling=50 separate");<br>setOption("BlackBackground", true);<br>run("Split Channels");<br>selectWindow("IMAGE_NAME.JPG (blue)");<br>close();<br>selectWindow(“IMAGE_NAME.JPG (red)");<br>close();<br>selectWindow("IMAGE_NAME.JPG (green)");<br>run("Convert to Mask"); |

Set scales before beginning to globally calibrate images and measurements. Use wand tool to select plant and measure area.

Table S4. All promoters that were used in this study. TATA boxes are highlighted in red, and USE are highlighted in green. Add coordinates of each promoter

| Species, promoter type and chromosome location. | Promoter sequence |
|---|---|
| *C. purpureus* U6<br>Chromosome 7<br>11559096-<br>11558782 | ACCAAATCCTAAACGATAGTGAAACCAAATTTAGACGAAACATATTCTCCAGCAGGT<br>CATATTCTACTAGTAGTAATCTTCGCAGTGTAAACTAATGCTCTCATCTCCTAAAAAC<br>GAGAAAAAAAGTAGTAAACTAAAGAAAGGAAGATGAGAAAGGACATTCTTCAGCA<br>AAAGGGAACACAACTTCAACGCAAAACATGAGGAAAAAGCAAAAGTATTCTGACA<br>CAACACCTCTATTTTAGGCTGACAAAAACCATAACGTTTGTCACGACGTATACCGGC<br>ATCCATATAACCACAGACGGAACCTAACCTATT<br>-23bp |
| *C. purpureus* U6<br>Chromosome 9<br>3874021-<br>3873707 | ATCTATGTCAAGGATGGACCACTCCTTTCCTTTCGCATCCACGAGGAGTTGCAAGTG<br>ACGTGCAACCAAAGCGCCATCTCAAGGTGAGAAATTCGAATGTGGTCACTGCTGCA<br>AGTTGATTTTGCTCCCATTTTGACATACACATGTTTCCTTCACTGCAAGGAGTCTAGT<br>AGGAATTTATGTTGCCATACTTTAGAATCATAGGACTATTTCGCAGCAGTACAGTAT<br>CAGGGATGCACTTGTGATCTGAAATCCCATAAGGATTGTGACTATAAACATAGCGA<br>ACTACTTAAGCTTGTGGTATCCTGCTTCCTTAT<br>-24 |
| *C. purpureus* U6<br>Chromosome 10<br>13131293-<br>13130980 | TTGCCATGACGAACTGTGGAATGAACCACCTAAGAGCTACTATCCTTGTTTATTTCC<br>ACTCGACTTATGTCATCTTGGAGTGGAAGTTAGAAGTAGTAGTTGGGTAGGCGACT<br>CACTGACCCAGAGGATACCTTACGCAAGGAAAGGGACCCTGCGTGGGCAAGTTCA<br>CATTTGCCCTGATGAAAAGCACACATACGGAGCATTGGGAAGGGCTGACATGGCAT<br>AGCATGGCATAGCATGGGAAGTATAACCCATATCGTTTGGGCAGGGCAGAGAGGC<br>GACCTATATA<br>-25 |
| *C. purpureus* U6<br>Chromosome 12<br>3954576-<br>3954890 | GCGAATGCCCAGAAGAAAGATGGCGGTTGTCGCACCTACATCATCGATGACAGTG<br>ATGCAGTGGGTGGTTGGGAAAGTGATGAGGTATGCCAAGACCTGCATACTGAGTC<br>AAACGACAGCCATGTCTGTTTTAAAGTTTTGCTGTTCTACCGGTACTCAAATCCCAG<br>AAATGATAAGTTGTTTCTTTTTCTGCAAGGTGCTGGAGTGGCATTAACTAAGATGGA<br>TACAGAGGATCGCATAATTGTCATATATACCCTAGGGTTAGCAACACTTTCGGAACC<br>GTAATATATATTACCCCAAACCCCTCATCCTTTC<br>-21 |
| P. *Patens* U6<br>Chromosome 1<br>5050307-<br>5050621 | GTCCATTGAAGCAGACGTGTTGCGACAGGTTAGCGACGATGGGTGTAGATGTGAT<br>GTGATGTGATGGTGTGGTTCTTCCACGGCGGCGTCCTTGCGGTGGCGGAGAAGGG<br>GATATCCCGAAGGAGCGGCAGCGGGAGAGCACAAGCAGAAAGGGTGCAGTGAGT<br>GAGTGGGTCCAGCTGGGTGGCTGGCCGAGTGGACGCGACCGGGTTTCGAGGGGG<br>CGGGGGAGAAAAGGGATGGAGCGAGGGATATAACCCACATGGAATGGAGGTGG<br>GTGTGAAGGCGGGTATATAGGAAGGTGGAGGACTTACAACCCAT<br>-25 |

| | |
|---|---|
| *C. purpureus* RpS5a Chromosome 3 19311369-19312666 | GGAGATAATAGAATAGTTGTTCCAAGGTGAACAAACATGTTAACAACTTACAATGT TCATTTGATTTTGGAGGAGTGAACTACGATCTTTTTATTAAAACCCACTACCTGGAT GTACATTAGATCATGTGCTCCAATAGAATGGTTGAATTTACATGTTGGCATTGAAGT ATACCTCATAGATATGAAGTATGTATGGAAAGATGGTATCAAGAAGATATGTATCT ATGTGTGAAAGTTGATTACTCTATGATACAGATGTATAGTTCTAAGATGTAGTAGTT GTGTACAATAGAGAGTAGTTGTATGTTGTAAGACCTTTAAATAGTAGTACTTTCAAC CAATACATGAGATGATGAAGAAGATTGACCCCTATAGGTTTTTGTGTTTTTGGAGCC AAAAGTGCCATAAACATACTCCCACTTCTCACCCTCCCTAGAAAAACCAAACTGGTT TAAAATAGATTTGGCCGGGTACGTTGATAGACCCCTAAGAATTGGAAAGGACCTTT CCTTGTACTTTACTCGTCATTTCTTCCACTTTTTCAAATTTATGAGCAGCCATAAGTCC GTCATTAAATAGTTACATTCAAACGCCTTGGTTTATAGTGGGTCATTTGAGCGAAAT ATTTGCATGCACCTATCAAAAAGTCTAGGGTCCAACACTGTTATTTTCATGCCAATCT GGTTTTTCGAGGGGGGTGAGACTTGTAGCAATGTAATGATTTCTAAGTTTAGACAA CCAATTGAGTCTTGAATCTTAGAGTCTTGTACCTACTCTATTGAAAGAAGTGTGTAT AAAGTTTGAACACAGTTCATTGTATAGATTGCCAAAGTTTAAGGTTTGAAGTCACTA TAACATGAGGATATTTGAACAATGTTAGGCTGTAATTAGTTGTTCTGTTTTAGGAAA CTAAACTAGTTTTTGGAAGATCGAATCTTATGTTCCAAGAAAAAGAAGAAGATATC GGTTTGAAGGAATGTTGAATTATTTTGGTCCAGCGGGCAGTGAAGAGAGTACAGTA GAGTTGTATTACCGCATTCCCGGGCACGCTCACAAACCAAGTTCCCTGCGGCGAAG CCCACGGGAACCATGCACCGTCAGATCGTACGGGCAGATCAGGGGCCGTCGATCT GTGACCGCGCCAGAGCACCTGGGTGCCCGGGAGAAAATTAGGGCGGGGGAGGGA GAAGGCCAAGACGGCCGAGAGAGCGCTAGCGGAGTCTCTTTCTCGACTCGCCATA GCGCCTGAGCAGCCTGCTCCCGCACGCGACGACGCCCCTCGCCCCTCCGCAACTCC ATC |

Table S5. Names of all moss isolates used for phenotypic evaluation of APT-KO mutants vs wildtype growth area.

| Isolate name | Shorthand name | Mutation |
|---|---|---|
| B150- APT#1#2-A | B150-A | Double cut 340bp del |
| B150-APT#1-B | B150-B | 9bp del MMEJ |
| B150-APT#1-C | B150-C | 22bp insert NHEJ |
| B190-APT#1#2-A | B190-A | Double cut MMEJ |
| B190-APT#1-B | B190-B | NHEJ 13bp Del |
| B190-APT#1-C | B190-C | MMEJ 10bp del |

Table S6. Number of RNA Polymerase III subunits found in male and female *Ceratodon purpureus* isolates. Highlighted in yellow are proteins found on male (V) or female (U) sex chromosomes.

| |
|---|
| CepurGG1.10G050400 - (1 of 1) KOG0260 - RNA polymerase II, large subunit |
| CepurGG1.10G098200 - (1 of 3) K03010 - DNA-directed RNA polymerase II subunit RPB2 (RPB2, POLR2B) |
| CepurGG1.10G102800 - (1 of 1) K03022 - DNA-directed RNA polymerase III subunit RPC8 (RPC8, POLR3H) |
| CepurGG1.10G148000 - (1 of 1) PF00623//PF04983//PF04997 - RNA polymerase Rpb1, domain 2 (RNA_pol_Rpb1_2) // RNA polymerase Rpb1, domain 3 (RNA_pol_Rpb1_3) // RNA polymerase Rpb1, domain 1 (RNA_pol_Rpb1_1) |
| CepurGG1.11G088500 - (1 of 1) KOG4168 - Predicted RNA polymerase III subunit C17 |
| CepurGG1.11G133600 - (1 of 1) PTHR12709:SF3 - DNA-DIRECTED RNA POLYMERASE IV SUBUNIT 7-RELATED |
| CepurGG1.12G027900 - (1 of 8) PF11523 - Protein of unknown function (DUF3223) (DUF3223) |
| CepurGG1.12G037200 - (1 of 1) K03024 - DNA-directed RNA polymerase III subunit RPC7 (RPC7, POLR3G) |
| CepurGG1.12G038200 - (1 of 1) K03013 - DNA-directed RNA polymerases I, II, and III subunit RPABC1 (RPB5, POLR2E) |
| CepurGG1.12G144400 - (1 of 1) PTHR33415//PTHR33415:SF3 - FAMILY NOT NAMED // EMB514 |
| CepurGG1.12G146300 - (1 of 1) K03023 - DNA-directed RNA polymerase III subunit RPC3 (RPC3, POLR3C) |
| CepurGG1.12G167400 - (1 of 2) K03021 - DNA-directed RNA polymerase III subunit RPC2 (RPC2, POLR3B) |
| CepurGG1.1G101300 - (1 of 2) K03020 - DNA-directed RNA polymerases I and III subunit RPAC2 (RPC19, POLR1D) |
| CepurGG1.1G126200 - (1 of 1) K03018 - DNA-directed RNA polymerase III subunit RPC1 (RPC1, POLR3A) |
| CepurGG1.1G137500 - (1 of 1) 2.7.7.48//2.7.7.6 - RNA-directed RNA polymerase / RNA nucleotidyltransferase (RNA-directed) // DNA-directed RNA polymerase / RNA polymerase III |
| CepurGG1.1G245900 - (1 of 2) K10908 - DNA-directed RNA polymerase, mitochondrial [EC:2.7.7.6] (POLRMT, RPO41) |
| CepurGG1.2G038700 - (1 of 2) K03008 - DNA-directed RNA polymerase II subunit RPB11 (RPB11, POLR2J) |
| CepurGG1.2G085300 - (1 of 2) PTHR19376//PTHR19376:SF37 - DNA-DIRECTED RNA POLYMERASE // DNA-DIRECTED RNA POLYMERASE II SUBUNIT RPB1 |
| CepurGG1.2G117200 - (1 of 2) PTHR19376//PTHR19376:SF37 - DNA-DIRECTED RNA POLYMERASE // DNA-DIRECTED RNA POLYMERASE II SUBUNIT RPB1 |
| CepurGG1.2G148700 - (1 of 3) KOG2351 - RNA polymerase II, fourth largest subunit |
| CepurGG1.2G157500 - (1 of 1) K02999 - DNA-directed RNA polymerase I subunit RPA1 (RPA1, POLR1A) |

CepurGG1.2G208800 - (1 of 1) K03011 - DNA-directed RNA polymerase II subunit RPB3 (RPB3, POLR2C)
CepurGG1.3G013100 - (1 of 1) PTHR11618:SF4 - TRANSCRIPTION FACTOR IIIB 90 KDA SUBUNIT
CepurGG1.3G091900 - (1 of 5) K03006 - DNA-directed RNA polymerase II subunit RPB1 (RPB1, POLR2A)
CepurGG1.3G097400 - (1 of 1) PTHR22504:SF0 - REPRESSOR OF RNA POLYMERASE III TRANSCRIPTION MAF1 HOMOLOG
CepurGG1.3G164800 - (1 of 2) K03012 - DNA-directed RNA polymerase II subunit RPB4 (RPB4, POLR2D)
CepurGG1.3G217100 - (1 of 1) K03000 - DNA-directed RNA polymerase I subunit RPA12 (RPA12, ZNRD1)
CepurGG1.3G222400 - (1 of 47) 2.7.7.6 - DNA-directed RNA polymerase / RNA polymerase III
CepurGG1.4G204000 - (1 of 2) PTHR19376//PTHR19376:SF11 - DNA-DIRECTED RNA POLYMERASE // DNA-DIRECTED RNA POLYMERASE I SUBUNIT RPA1
CepurGG1.5G090200 - (1 of 2) PTHR12949:SF0 - DNA-DIRECTED RNA POLYMERASE III SUBUNIT RPC3
CepurGG1.5G167800 - (1 of 2) K03020 - DNA-directed RNA polymerases I and III subunit RPAC2 (RPC19, POLR1D)
CepurGG1.6G036600 - (1 of 2) PTHR12709//PTHR12709:SF1 - DNA-DIRECTED RNA POLYMERASE II, III // DNA-DIRECTED RNA POLYMERASE III SUBUNIT RPC8
CepurGG1.6G070000 - (1 of 1) K03007 - DNA-directed RNA polymerases I, II, and III subunit RPABC5 (RPB10, POLR2L)
CepurGG1.6G205100 - (1 of 1) K14721 - DNA-directed RNA polymerase III subunit RPC5 (RPC5, POLR3E)
CepurGG1.7G047100 - (1 of 3) KOG0261 - RNA polymerase III, large subunit
CepurGG1.7G047400 - (1 of 7) PTHR19376//PTHR19376:SF40 - DNA-DIRECTED RNA POLYMERASE // DNA-DIRECTED RNA POLYMERASE IV SUBUNIT 1
CepurGG1.7G050200 - (1 of 1) K03040 - DNA-directed RNA polymerase subunit alpha (rpoA)
CepurGG1.7G056800 - (1 of 3) KOG0261 - RNA polymerase III, large subunit
CepurGG1.7G058800 - (1 of 2) PTHR17204//PTHR17204:SF29 - PRE-MRNA PROCESSING PROTEIN PRP39-RELATED // SUBFAMILY NOT NAMED
CepurGG1.7G076100 - (1 of 1) K16251 - DNA-directed RNA polymerase V subunit 1 [EC:2.7.7.6] (NRPE1)
CepurGG1.7G077400 - (1 of 7) PTHR19376//PTHR19376:SF40 - DNA-DIRECTED RNA POLYMERASE // DNA-DIRECTED RNA POLYMERASE IV SUBUNIT 1
CepurGG1.7G086300 - (1 of 3) K03010 - DNA-directed RNA polymerase II subunit RPB2 (RPB2, POLR2B)
CepurGG1.7G094000 - (1 of 1) K03019 - DNA-directed RNA polymerase III subunit RPC11 (RPC11, POLR3K)
CepurGG1.7G120400 - (1 of 2) K10908 - DNA-directed RNA polymerase, mitochondrial [EC:2.7.7.6] (POLRMT, RPO41)

CepurGG1.7G160100 - (1 of 2) K03012 - DNA-directed RNA polymerase II subunit RPB4 (RPB4, POLR2D)
CepurGG1.8G012600 - (1 of 1) PTHR12709//PTHR12709:SF4 - DNA-DIRECTED RNA POLYMERASE II, III // DNA-DIRECTED RNA POLYMERASE II SUBUNIT RPB7
CepurGG1.8G052100 - (1 of 47) 2.7.7.6 - DNA-directed RNA polymerase / RNA polymerase III
CepurGG1.8G071300 - (1 of 1) K15198 - transcription factor TFIIIB component B'' (BDP1, TFC5)
CepurGG1.8G100700 - (1 of 1) K03017 - DNA-directed RNA polymerase II subunit RPB9 (RPB9, POLR2I)
CepurGG1.8G174200 - (1 of 1) K03025 - DNA-directed RNA polymerase III subunit RPC6 (RPC6, POLR3F)
CepurGG1.9G031500 - (1 of 1) K03014 - DNA-directed RNA polymerases I, II, and III subunit RPABC2 (RPB6, POLR2F)
CepurGG1.9G160100 - (1 of 1) K03026 - DNA-directed RNA polymerase III subunit RPC4 (RPC4, POLR3D)
CepurGG1.9G174800 - (1 of 2) PTHR13408//PTHR13408:SF0 - DNA-DIRECTED RNA POLYMERASE III // DNA-DIRECTED RNA POLYMERASE III SUBUNIT RPC4
CepurGG1.N011700 - (1 of 2) K03008 - DNA-directed RNA polymerase II subunit RPB11 (RPB11, POLR2J)
CepurGG1.UG179700 - (1 of 2) K03021 - DNA-directed RNA polymerase III subunit RPC2 (RPC2, POLR3B)
CepurGG1.UG204100 - (1 of 1) K03002 - DNA-directed RNA polymerase I subunit RPA2 (RPA2, POLR1B)
CepurGG1.UG292500 - (1 of 1) PTHR20856:SF7 - DNA-DIRECTED RNA POLYMERASE II SUBUNIT RPB2
CepurGG1.UG319900 - (1 of 2) K03015 - DNA-directed RNA polymerase II subunit RPB7 (RPB7, POLR2G)
CepurGG1.UG335800 - (1 of 1) K03027 - DNA-directed RNA polymerases I and III subunit RPAC1 (RPC40, POLR1C)
CepurR40.10G053800 - (1 of 2) KOG0260 - RNA polymerase II, large subunit
CepurR40.10G100300 - (1 of 3) K03010 - DNA-directed RNA polymerase II subunit RPB2 (RPB2, POLR2B)
CepurR40.10G106000 - (1 of 1) K03022 - DNA-directed RNA polymerase III subunit RPC8 (RPC8, POLR3H)
CepurR40.10G153400 - (1 of 1) PF00623//PF04983//PF04997 - RNA polymerase Rpb1, domain 2 (RNA_pol_Rpb1_2) // RNA polymerase Rpb1, domain 3 (RNA_pol_Rpb1_3) // RNA polymerase Rpb1, domain 1 (RNA_pol_Rpb1_1)
CepurR40.11G027900 - (1 of 1) 2.7.7.48//2.7.7.6 - RNA-directed RNA polymerase / RNA nucleotidyltransferase (RNA-directed) // DNA-directed RNA polymerase / RNA polymerase III
CepurR40.11G031300 - (1 of 2) KOG3233 - RNA polymerase III, subunit C34
CepurR40.11G091500 - (1 of 1) KOG4168 - Predicted RNA polymerase III subunit C17

CepurR40.11G137100 - (1 of 1) PTHR12709:SF3 - DNA-DIRECTED RNA POLYMERASE IV SUBUNIT 7-RELATED
CepurR40.11G154400 - (1 of 4) PTHR19376//PTHR19376:SF11 - DNA-DIRECTED RNA POLYMERASE // DNA-DIRECTED RNA POLYMERASE I SUBUNIT RPA1
CepurR40.12G028000 - (1 of 7) PF11523 - Protein of unknown function (DUF3223) (DUF3223)
CepurR40.12G036600 - (1 of 2) K03024 - DNA-directed RNA polymerase III subunit RPC7 (RPC7, POLR3G)
CepurR40.12G037500 - (1 of 1) K03013 - DNA-directed RNA polymerases I, II, and III subunit RPABC1 (RPB5, POLR2E)
CepurR40.12G140800 - (1 of 1) PTHR33415//PTHR33415:SF3 - FAMILY NOT NAMED // EMB514
CepurR40.12G142600 - (1 of 1) K03023 - DNA-directed RNA polymerase III subunit RPC3 (RPC3, POLR3C)
CepurR40.12G165000 - (1 of 2) K03021 - DNA-directed RNA polymerase III subunit RPC2 (RPC2, POLR3B)
CepurR40.12G173900 - (1 of 7) PF11523 - Protein of unknown function (DUF3223) (DUF3223)
CepurR40.1G112300 - (1 of 2) K03020 - DNA-directed RNA polymerases I and III subunit RPAC2 (RPC19, POLR1D)
CepurR40.1G130500 - (1 of 1) K03018 - DNA-directed RNA polymerase III subunit RPC1 (RPC1, POLR3A)
CepurR40.1G134700 - (1 of 4) PTHR19376//PTHR19376:SF11 - DNA-DIRECTED RNA POLYMERASE // DNA-DIRECTED RNA POLYMERASE I SUBUNIT RPA1
CepurR40.1G251500 - (1 of 2) K10908 - DNA-directed RNA polymerase, mitochondrial [EC:2.7.7.6] (POLRMT, RPO41)
CepurR40.2G039500 - (1 of 2) K03008 - DNA-directed RNA polymerase II subunit RPB11 (RPB11, POLR2J)
CepurR40.2G085700 - (1 of 2) PTHR19376//PTHR19376:SF37 - DNA-DIRECTED RNA POLYMERASE // DNA-DIRECTED RNA POLYMERASE II SUBUNIT RPB1
CepurR40.2G137000 - (1 of 2) PTHR19376//PTHR19376:SF37 - DNA-DIRECTED RNA POLYMERASE // DNA-DIRECTED RNA POLYMERASE II SUBUNIT RPB1
CepurR40.2G176100 - (1 of 2) K03008 - DNA-directed RNA polymerase II subunit RPB11 (RPB11, POLR2J)
CepurR40.2G185400 - (1 of 1) K02999 - DNA-directed RNA polymerase I subunit RPA1 (RPA1, POLR1A)
CepurR40.2G218100 - (1 of 4) KOG2351 - RNA polymerase II, fourth largest subunit
CepurR40.2G254300 - (1 of 1) K03011 - DNA-directed RNA polymerase II subunit RPB3 (RPB3, POLR2C)
CepurR40.3G013000 - (1 of 1) PTHR11618:SF4 - TRANSCRIPTION FACTOR IIIB 90 KDA SUBUNIT
CepurR40.3G094300 - (1 of 3) PTHR22504:SF0 - REPRESSOR OF RNA POLYMERASE III TRANSCRIPTION MAF1 HOMOLOG

CepurR40.3G165400 - (1 of 3) K03012 - DNA-directed RNA polymerase II subunit RPB4 (RPB4, POLR2D)
CepurR40.3G216200 - (1 of 1) K03000 - DNA-directed RNA polymerase I subunit RPA12 (RPA12, ZNRD1)
CepurR40.3G220700 - (1 of 55) 2.7.7.6 - DNA-directed RNA polymerase / RNA polymerase III
CepurR40.5G074800 - (1 of 2) KOG2587 - RNA polymerase III (C) subunit
CepurR40.5G174700 - (1 of 2) K03020 - DNA-directed RNA polymerases I and III subunit RPAC2 (RPC19, POLR1D)
CepurR40.6G074600 - (1 of 1) K03007 - DNA-directed RNA polymerases I, II, and III subunit RPABC5 (RPB10, POLR2L)
CepurR40.6G214200 - (1 of 1) K14721 - DNA-directed RNA polymerase III subunit RPC5 (RPC5, POLR3E)
CepurR40.7G050700 - (1 of 2) KOG0261 - RNA polymerase III, large subunit
CepurR40.7G050800 - (1 of 6) PTHR19376//PTHR19376:SF40 - DNA-DIRECTED RNA POLYMERASE // DNA-DIRECTED RNA POLYMERASE IV SUBUNIT 1
CepurR40.7G053200 - (1 of 1) K03040 - DNA-directed RNA polymerase subunit alpha (rpoA)
CepurR40.7G059400 - (1 of 2) KOG0260 - RNA polymerase II, large subunit
CepurR40.7G068200 - (1 of 1) K16251 - DNA-directed RNA polymerase V subunit 1 [EC:2.7.7.6] (NRPE1)
CepurR40.7G093600 - (1 of 3) K03010 - DNA-directed RNA polymerase II subunit RPB2 (RPB2, POLR2B)
CepurR40.7G100400 - (1 of 1) K03019 - DNA-directed RNA polymerase III subunit RPC11 (RPC11, POLR3K)
CepurR40.7G120400 - (1 of 4) PTHR19376//PTHR19376:SF11 - DNA-DIRECTED RNA POLYMERASE // DNA-DIRECTED RNA POLYMERASE I SUBUNIT RPA1
CepurR40.7G156500 - (1 of 2) K10908 - DNA-directed RNA polymerase, mitochondrial [EC:2.7.7.6] (POLRMT, RPO41)
CepurR40.8G013300 - (1 of 1) PTHR12709//PTHR12709:SF4 - DNA-DIRECTED RNA POLYMERASE II, III // DNA-DIRECTED RNA POLYMERASE II SUBUNIT RPB7
CepurR40.8G054200 - (1 of 55) 2.7.7.6 - DNA-directed RNA polymerase / RNA polymerase III
CepurR40.8G076100 - (1 of 1) K15198 - transcription factor TFIIIB component B'' (BDP1, TFC5)
CepurR40.8G105100 - (1 of 1) K03017 - DNA-directed RNA polymerase II subunit RPB9 (RPB9, POLR2I)
CepurR40.8G179300 - (1 of 1) K03025 - DNA-directed RNA polymerase III subunit RPC6 (RPC6, POLR3F)
CepurR40.9G029500 - (1 of 1) K03014 - DNA-directed RNA polymerases I, II, and III subunit RPABC2 (RPB6, POLR2F)
CepurR40.9G064900 - (1 of 1) PTHR13408//PTHR13408:SF3 - DNA-DIRECTED RNA POLYMERASE III // SUBFAMILY NOT NAMED
CepurR40.9G072000 - (1 of 1) PF00562//PF04560 - RNA polymerase Rpb2, domain 6 (RNA_pol_Rpb2_6) // RNA polymerase Rpb2, domain 7 (RNA_pol_Rpb2_7)
CepurR40.9G072100 - (1 of 3) PTHR20856:SF7 - DNA-DIRECTED RNA POLYMERASE II SUBUNIT RPB2

CepurR40.9G132600 - (1 of 1) K03026 - DNA-directed RNA polymerase III subunit RPC4 (RPC4, POLR3D)
CepurR40.N000200 - (1 of 3) PTHR22504:SF0 - REPRESSOR OF RNA POLYMERASE III TRANSCRIPTION MAF1 HOMOLOG
CepurR40.N001400 - (1 of 2) K03024 - DNA-directed RNA polymerase III subunit RPC7 (RPC7, POLR3G)
CepurR40.N002400 - (1 of 2) PTHR10773 - DNA-DIRECTED RNA POLYMERASES I, II, AND III SUBUNIT RPABC2
CepurR40.N009700 - (1 of 3) K03012 - DNA-directed RNA polymerase II subunit RPB4 (RPB4, POLR2D)
CepurR40.N014500 - (1 of 3) K03012 - DNA-directed RNA polymerase II subunit RPB4 (RPB4, POLR2D)
CepurR40.N045400 - (1 of 3) PTHR22504:SF0 - REPRESSOR OF RNA POLYMERASE III TRANSCRIPTION MAF1 HOMOLOG
CepurR40.VG130100 - (1 of 2) K03015 - DNA-directed RNA polymerase II subunit RPB7 (RPB7, POLR2G)
CepurR40.VG151700 - (1 of 3) K03010 - DNA-directed RNA polymerase II subunit RPB2 (RPB2, POLR2B)
CepurR40.VG178800 - (1 of 5) PTHR20856//PTHR20856:SF8 - DNA-DIRECTED RNA POLYMERASE I SUBUNIT 2 // DNA-DIRECTED RNA POLYMERASE III SUBUNIT RPC2
CepurR40.VG178900 - (1 of 5) PTHR20856//PTHR20856:SF8 - DNA-DIRECTED RNA POLYMERASE I SUBUNIT 2 // DNA-DIRECTED RNA POLYMERASE III SUBUNIT RPC2
CepurR40.VG179200 - (1 of 5) PTHR20856//PTHR20856:SF8 - DNA-DIRECTED RNA POLYMERASE I SUBUNIT 2 // DNA-DIRECTED RNA POLYMERASE III SUBUNIT RPC2
CepurR40.VG179300 - (1 of 55) 2.7.7.6 - DNA-directed RNA polymerase / RNA polymerase III
CepurR40.VG255100 - (1 of 1) K03002 - DNA-directed RNA polymerase I subunit RPA2 (RPA2, POLR1B)
CepurR40.VG314600 - (1 of 2) K03021 - DNA-directed RNA polymerase III subunit RPC2 (RPC2, POLR3B)
CepurR40.VG336200 - (1 of 1) K03027 - DNA-directed RNA polymerases I and III subunit RPAC1 (RPC40, POLR1C)

**Datasets**

Data S1.

All Sanger sequencing data of CpAPT-KO mutants and CpLand inserts.
